# Music-to My Brain: Modulation of Reward System Activity via Musical Neurofeedback

**DOI:** 10.64898/2026.05.04.718856

**Authors:** N. Singer, A. Rabinowitz, Y. Koryto-Blumen, Y. Hamrani, M. Farres-Franch, M. Doron, N. Dunsky, G. Gurevitch, T. Hendler, A. Dagher, R.J Zatorre

## Abstract

Music robustly engages the human reward system, yet whether this engagement can be harnessed for volitional self-neuromodulation remains unknown. We developed a musical neurofeedback approach that enables individuals to control a validated, fMRI-informed EEG marker of ventral striatal activity. Personalized pleasurable music served as feedback, becoming increasingly rewarding through acoustic manipulation as regulation improved, thereby creating a positive feedback loop.

Across three double-blind, sham-controlled studies (N=80; two with repeated training), contingent neurofeedback enabled participants to upregulate this EEG signal reflecting ventral striatal activity; in studies with repeated training, this learning generalized to no-feedback contexts.

In the neurofeedback (but not sham) group, regulation success correlated with self-reported hedonic capacity, indicating behavioral relevance. Pre-post fMRI further showed that improvements in ventral striatal BOLD self-regulation were associated with EEG-based regulation performance, supporting the EEG measure as a marker of ventral striatal modulation. Mechanistically, neurofeedback training enhanced functional connectivity between right auditory cortex and ventral striatum during listening to trained music, with stronger effects in the neurofeedback than in the sham group, demonstrating experience-dependent modification of auditory-reward pathways.

Together, these findings reveal a mechanism for music-based reward self-regulation and offer a potential scalable, personalized approach for targeting reward dysfunction such as anhedonia.

## Introduction

Music is a universal human experience with profound effects on emotion and well-being. One remarkable example is music’s capacity to induce intense pleasure, a process closely tied to engagement of the brain’s reward circuitry^1^. Musical pleasure reliably engages mesolimbic dopaminergic circuits, particularly the ventral striatum (VS)— a key node of the reward circuitry^1–5^ that has long been recognized as essential for processing biological reward in both animals and humans^6–8^. Musical pleasure is also associated with phasic dopamine release in the striatum^9^ and enhanced functional connectivity between auditory and reward regions^2,3,10^. This auditory-reward interaction is theorized as the mechanistic pathway for music’s hedonic value^11^. Beyond its aesthetic appeal, music is widely used therapeutically to regulate emotions and reduce stress, anxiety, and depression^12–19^. Together, these findings raise a fundamental question: can music’s naturally occurring reward-circuitry engagement be harnessed for volitional control of reward-related brain activity?

Neurofeedback, a method enabling individuals to learn volitional control of brain activity using real-time feedback^20,21^, offers a direct way to test whether pleasurable music can serve as an effective medium for reward circuit self-regulation. Demonstrating such regulation would reveal whether auditory-reward pathways can be harnessed for volitional training and establish a potential neural-guided approach for targeting symptoms associated with mesolimbic dysregulation, such as anhedonia and apathy^22,23^.

Previous neurofeedback studies have demonstrated that individuals can self-regulate activation in major nodes of reward circuitry such as the VS and Ventral Tegmental Area (VTA) using real-time fMRI with visual feedback interfaces^24–30^. Yet, these approaches have faced practical limitations and often failed to show transfer to behavior or regulation without feedback^25,28,29^. First, fMRI-based neurofeedback is constrained by cost, immobility, and the unnatural scanning environment, restricting its applicability for long-term training in naturalistic settings^31,32^. Second, these studies have typically employed neutral visual feedback (e.g., bar graphs) rather than feedback that engages the target system or process directly, which may limit regulation efficacy^21,33^.

Here, we present a novel musical neurofeedback approach that directly exploits the rewarding value of music to enable self-modulation of reward-related processing while addressing both of these limitations of current neurofeedback approaches. Specifically, we leverage a recently validated fMRI-informed EEG model, the VS-Electrical FingerPrint (VS-EFP), which tracks VS blood-oxygen-level-dependent (BOLD)-related activation. This EEG model was trained on simultaneous EEG/fMRI data collected during pleasurable music listening³⁵ and provides a scalable (accessible and affordable) alternative to fMRI while maintaining target specificity. We use VS-EFP signal modulations to manipulate pleasurable, self-selected music in real time, such that its acoustic quality, and thus reward value, scales with the user’s success in upregulating the VS-EFP signal. In this way, music serves both as feedback informing regulation success^34^ and as an engaging input to the reward system in a personalized manner^1,9^, creating a closed auditory-reward loop that may enhance learning efficacy^21^.

We tested this approach across three sham-controlled, double-blind studies with healthy volunteers: two proof-of-concept experiments and a preregistered replication with a larger sample (Fig. 1). Study 1 (N = 20) tested whether individuals could upregulate their VS-EFP signal with musical neurofeedback driven by their own VS-EFP modulation (test group) relative to sham feedback driven by another participant’s signal (sham group). Study 2 (N = 20) examined whether this neuromodulatory skill could be sustained and improved through six repeated musical neurofeedback sessions, assessed learning dynamics and generalization to no-feedback contexts (transfer capacity), and evaluated neurobehavioral outcomes including reward-related behavior and VS-BOLD regulation measured with pre-post fMRI. Study 3 (N = 40; preregistered) replicated key findings from Study 2 in a larger sample and explored the underlying neural mechanisms, specifically testing for training-induced changes in functional connectivity between auditory cortex and VS during music listening, as proposed in current cognitive neuroscience models of musical pleasure^11^.

**Figure 1.**
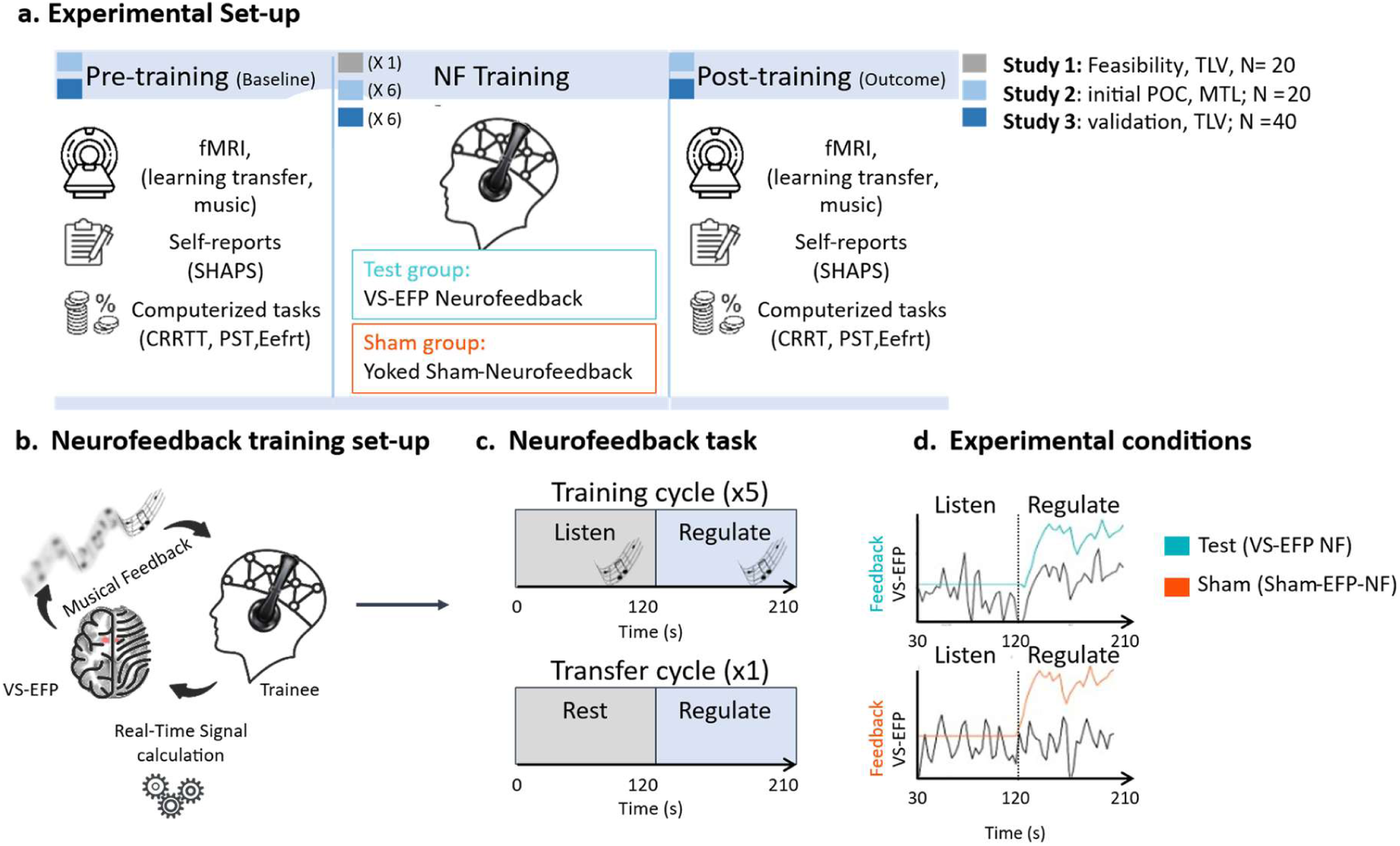
Experimental design and VS-EFP neurofeedback protocol. **(a)** *Study design*. Study 1: single-session feasibility. Studies 2-3: six training sessions over 2-3 weeks with pre-post assessments including fMRI and behavioral tasks. **(b)** *Neurofeedback training set-up*. Successful VS-EFP upregulation improves sound quality of participant-selected music in real-time. **(c)** *Neurofeedback task*. Five cycles alternating between passive listening (baseline) and regulation (participants find mental strategies to improve sound quality through VS-EFP upregulation). **(d)** *Experimental conditions*. Participants randomized to test (contingent VS-EFP feedback; nS1=10, nS2=10, nS3=20) or sham (yoked feedback; nS1=10, nS2=10, nS3=20). Example traces show a test participant receiving contingent feedback based on their own VS-EFP signal and their yoked sham participant receiving feedback based on the test participant’s signal. Study 3: half of sham group received reversed feedback. Abbreviations: VS = ventral striatum; EFP = electrical fingerprint; SHAPS = Snaith-Hamilton Pleasure Scale; EEfR = Effort Expenditure for Rewards Task; PST = Probabilistic Selection Task; CRRT = Cued Reinforcement Reaction Task.

We hypothesized that (1) VS-EFP neuromodulation capacity would be higher in the test group, receiving real-time neurofeedback, compared to the sham group (Study 1); (2) repeated training over sessions would enhance VS-EFP upregulation capacity and enable transfer of this capacity to no-feedback contexts (Studies 2-3); (3) Training success would correlate with self-reported hedonic capacity (defined as the individual’s ability to experience consummatory pleasure across various life domains) and produce changes in reward-related behaviors (Studies 2-3); (4) VS-EFP upregulation during training would correlate with improvements in VS-BOLD upregulation in the absence of feedback (Studies 2-3); and (5) musical neurofeedback would enhance functional connectivity between auditory cortex and VS (Study 3).

## Results

### A. Musical neurofeedback enables volitional modulation of the VS-EFP signal (Study 1)

In this initial feasibility study, we tested whether individuals could volitionally modulate the VS-EFP signal using real-time musical neurofeedback, and whether this capacity relates to hedonic capacity (the ability to experience pleasure, assessed via the Snaith-Hamilton Pleasure Scale; SHAPS^35^). Twenty healthy individuals completed one neurofeedback session attempting to modulate their VS-EFP signal using feedback from self-selected music, where music volume increased proportionally with VS-EFP upregulation, such that successful upregulation was reinforced by louder, more salient music. Participants were randomly assigned to test (n = 10; real-time VS-EFP feedback) or sham (n = 10; yoked feedback from another participant) groups.

We tested for neurofeedback effects on VS-EFP self-neuromodulation capacity (defined as the mean VS-EFP power difference between ‘regulate’ and passive ‘listen’ conditions; Fig. 2a)^36^. Descriptively, both groups significantly modulated their VS-EFP activity relative to baseline (One Sample Wilcoxon signed rank test; test: W = 54, p = 0.002; sham: W = 49, p = 0.01, one-tailed). Critically, a linear mixed model, with group (test vs sham) as a fixed effect, cycle (regulation runs 1 to 5) as a nuisance covariate, and random intercepts for participants, revealed a significant group effect (*β* = 0.04; 95% CI [0.01, 0.07]; *F*(1,17.49) = 5.2, *p* = 0.04, η² = 0.23), confirming greater VS-EFP upregulation with a contingent real-time musical neurofeedback.

**Figure 2.**
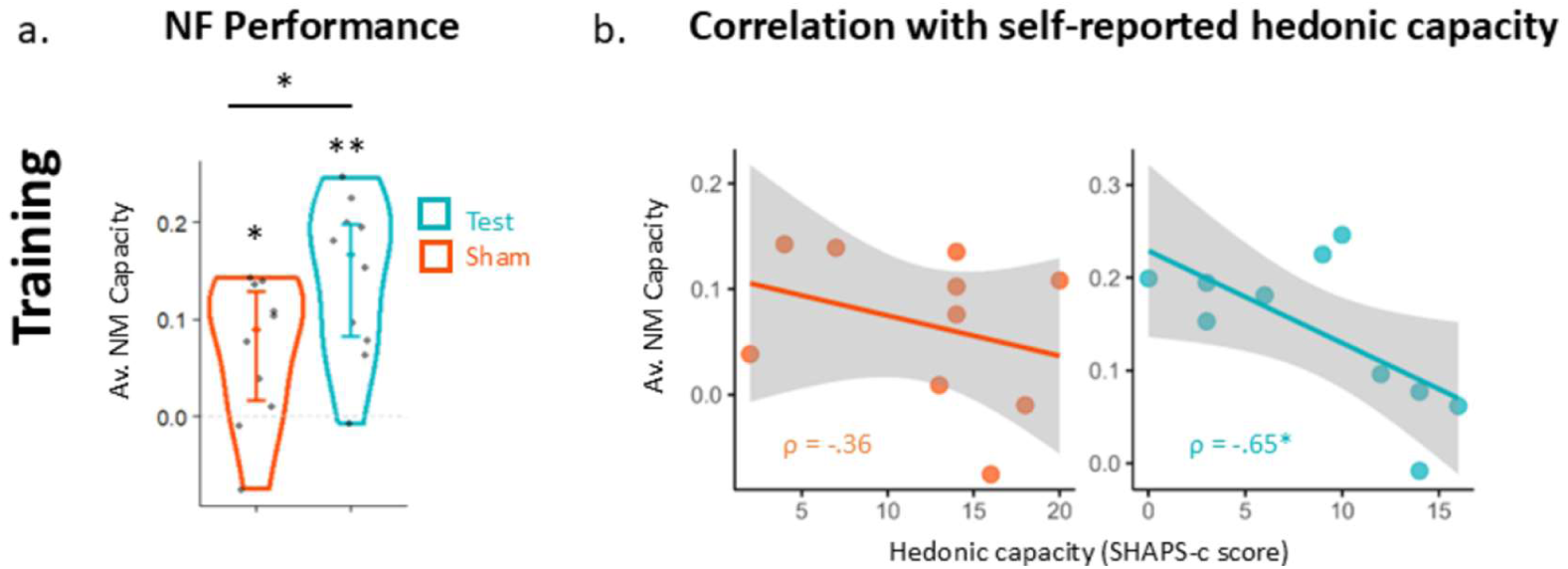
Study 1 results: A sham-controlled NF-EFP pilot to examine the feasibility of modulating the VS-EFP via neurofeedback (NF). **(a)** *NF neuromodulation capacity*, defined as the mean VS-EFP power difference between ‘regulate’ and passive ‘listen’ conditions, averaged across cycles for the test (cyan-blue, n=10) and the sham group (orange, n=10). Presented as violin plots overlaid with individual data points. **(b)** Correlations between NF neuromodulation capacity and self-reported hedonic capacity within the test (cyan-blue) and control (orang) group, with regression lines and 95% confidence intervals. Error bars represent the standard error of the mean (SEM). Abbreviations: NM = neuromodulation; VS = ventral striatum, EFP = electrical fingerprint. * p < 0.05; ** p < .01; one-sided hypothesis.

To assess behavioral relevance, we examined associations with individual differences in reward processing. VS-EFP upregulation during training correlated with baseline hedonic capacity (assessed via the SHAPS questionnaire, where lower scores indicate greater ability to experience pleasure) in the test group (*ρ* = -.65, *p* = .02, one-sided; Fig. 2b), but not the sham group (*ρ* = -.36, *p* = 0.15, one-sided). Thus, participants with higher baseline hedonic capacity better learned to upregulate their VS-EFP with neurofeedback.

### B. Repeated VS-EFP neurofeedback training: learning and behavioral outcomes (Study 2)

Having established that the VS-EFP signal can be modulated through musical neurofeedback and the relevance of its modulation to hedonic capacity (Study 1), we next tested whether this regulatory capacity could be sustained and improved through repeated training, and characterized associated neurobehavioral outcomes. Twenty individuals were randomly assigned to test (*n* = 10; VS-EFP neurofeedback) or sham (*n* = 10; yoked sham neurofeedback) groups, completing six training sessions with pre-and post-training assessments. We optimized the protocol by shortening the regulation phase to 90 seconds. Critically, we implemented a novel feedback scheme designed to modulate music’s reward value in real time — shifting from distorted, heavily filtered audio toward the participant’s preferred, high-fidelity version with greater VS-EFP upregulation relative to baseline — thereby creating a positive feedback loop in which successful self-regulation was reinforced by increasingly pleasurable music.

B.1. Real-time VS-EFP upregulation is greater with contingent than sham feedback. We first examined whether individuals could learn to self-modulate the VS-EFP signal across multiple sessions. Descriptively, one-sample Wilcoxon signed-rank tests per session revealed that the test group, but not the sham group, demonstrated significant neuromodulation capacity in the majority of sessions (test group: *q*(FDR) < .05 for sessions 1-3 and 6; *q*(FDR) = 0.06 for session 5; sham group: *q*(FDR) > 0.07 for all sessions, one-sided; Fig. 3a). However, direct between-group comparisons using linear mixed-effects modeling across training cycles and sessions revealed no significant group-by-session interaction (*F(5, 512.87)* = 1.04; p = 0.39; *β (linear)* = 0.00; 95% CI [- 0.02, 0.02]), nor main effects of group (*F(1, 9.58)* = 0.24; *p* =0.63; *β* = 0.01; 95% CI [- 0.02, 0.03]) or session (*F(5, 512.87)* = 0.64; *p* =0.67; *β (linear)* = -0.00; 95% CI [-0.01, 0.02]). This lack of group-level differentiation despite test group effects may reflect greater variability in the sham group (SDsham = 0.077, SDtest = 0.04). We hypothesized this may have arisen from incidental alignment between participants’ own VS-EFP signals and their yoked sham feedback — when a participant’s neural dynamics coincidentally matched another’s feedback, this partial contingency may have reinforced learning. Supporting this, the correlation between participants’ own VS-EFP signals and received feedback predicted neuromodulation capacity in the sham group (*F*(1, 278.25) = 9.04, *p* = 0.003; Fig. 3b) and across both groups (*F*(1, 21.47) = 7.23, *p* = 0.01). These findings suggest that signal-feedback contingency was associated with neuromodulation capacity, even when contingency was partial as in the sham group.

**Figure 3:**
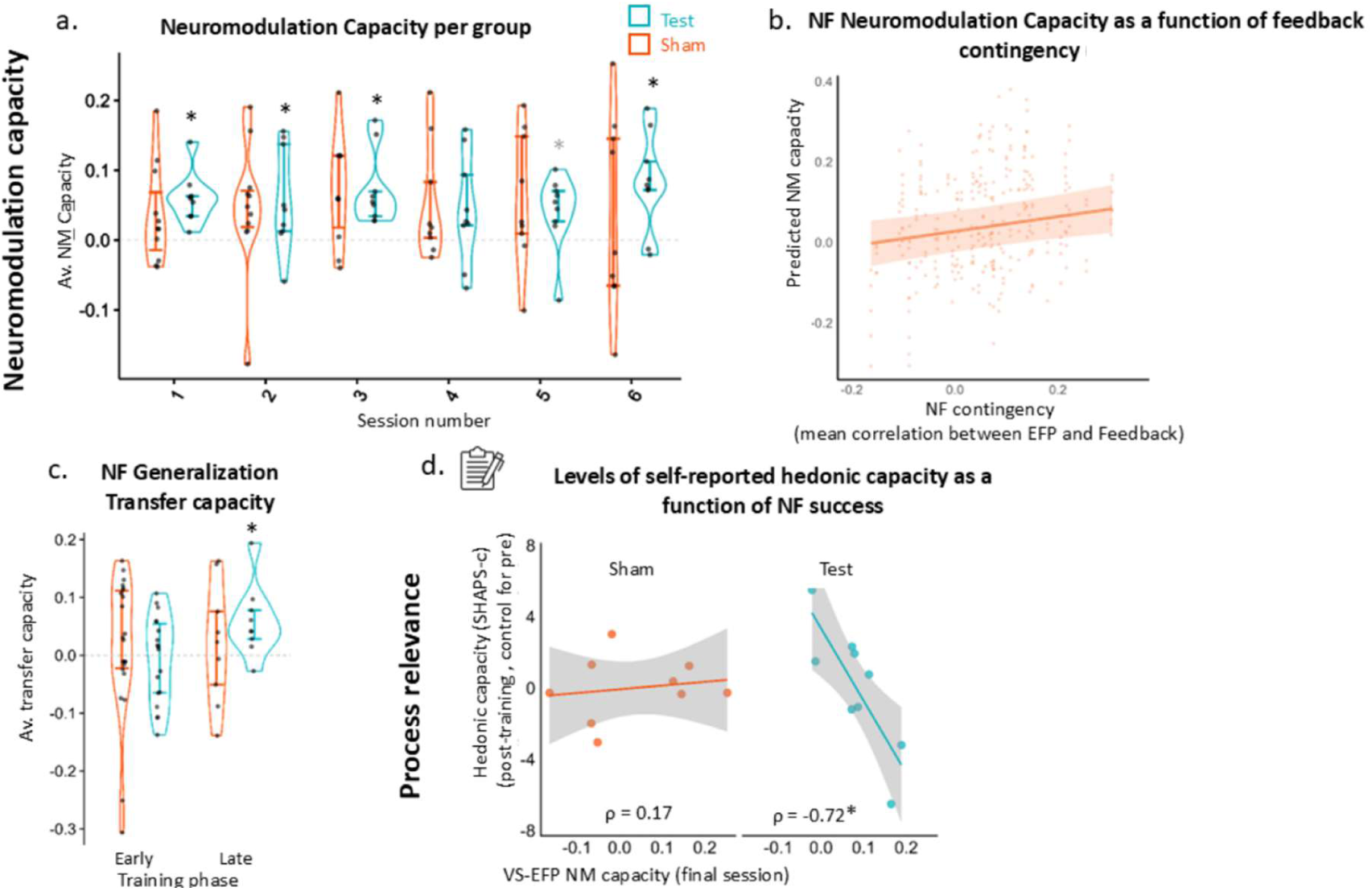
Study 2 results: Feasibility of learning during repeated VS-EFP NF sessions: Capacity, Generalization, and Associations with Hedonic Capacity. (**a**) *Neuromodulation (NM) capacity,* across cycles for each of the six sessions, shown separately for the test group (*n* = 9) and the sham group (*n* = 9) using violin plots with individual data points. NM capacity is defined as the mean VS-EFP power difference between ‘regulate’ and passive ‘listen’ conditions (**b**) *NF contingency predicts NF capacity in the sham group*. Predicted relationship between NF capacity and the extent of contingency between the VS-EFP signal and the feedback (mean correlation) in the sham group, as estimated from a linear mixed-effects model. (**c**) *Learning* - *Generalization*: transfer capacity averaged across early (sessions 2-3) and late (sessions 4-6) phases of training per group, depicted using violin plots with individual data points. (**d**) *Process relevance*: correlation between NF capacity during the last NF training session and post-training hedonic capacity (controlling for baseline hedonic capacity), plotted per group with regression lines and 95% confidence intervals. Error bars represent the standard error of the mean (SEM). Asterisks indicate significant differences (\**p* < .05), with asterisks above violins indicating differences from zero (p < .05, FDR corrected).

B.2. Transfer capacity improves from early to late training in the neurofeedback group. We next examined transfer capacity — the ability to volitionally modulate the VS-EFP signal in the absence of real-time feedback, measured during a dedicated transfer cycle at the end of each training session — as an indicator of learning generalization. Transfer capacity was quantified as the difference between regulation and baseline VS-EFP [Δ regulate - baseline] during the transfer cycles, assessed across early and late training phases. We first examined whether each group achieved above-baseline transfer capacity at each phase. Descriptively, VS-EFP neurofeedback, but not sham, was associated with enhanced transfer capacity during late training (late phase: test: *q*(FDR) = 0.02; sham: *q*(FDR) = 0.33; early phase: test: *q*(FDR) = 0.9; sham: *q*(FDR) = 0.31, one-sided; Fig. 3c). Testing specificity, i.e., whether groups differed in their transfer trajectories, linear mixed models revealed a group-by-training phase interaction (*F(1, 69.24)* = 7.02; *p* = 0.01; *β* = 0.03, CI [0.01, 0.04], η² = 0.09). Post-hoc analysis showed greater transfer capacity in the test group during late versus early training (*t*(65.72) = 2.47, *p* = 0.02), indicating improved transfer capacity from early to late training phases in the test group (Fig 3c). There was no main effect for group (*F(1, 18.76)* = 0.06, p = 0.80) or training phase (*F(1, 67.12)* = 2.09, *p* = 0.15).

B.3. Neurofeedback training is associated with post-training hedonic capacity and improvement in reward-driven behavior. Having established that VS-EFP can be modulated and learned through neurofeedback, we next examined whether training was associated with reward-related behavior.

#### Self-reported hedonic capacity

We examined whether neurofeedback-induced neuromodulation was associated with hedonic capacity, measured via SHAPS before and after the six-session training period. In the test group, greater neuromodulation capacity during the final session was significantly correlated with post-training hedonic capacity, controlling for baseline (*ρ* = -0.72, *p* = 0.02, one-sided; Fig. 3d), whereas no such relationship was observed in the sham group (*ρ* = 0.17, *p* = 0.67).

#### Reward Behavior

To examine behavioral relevance beyond self-report, we conducted exploratory analyses on three computerized tasks probing reward processing, administered before and after training: the Probabilistic Selection Task (PST)^37^, the Effort Expenditure for Rewards Task (EEfRT)^38^, and the Cued Reinforcement Reaction Task (CRRT)^39,40^, a modified version of the Monetary Incentive Delay task^41^. Here we focus on the CRRT, which showed replicable effects across studies 2 and 3.

In the CRRT, participants responded to visual targets following anticipatory cues indicating reward magnitude potential (high, low, or none), with reaction time (RT) proportionally determining the monetary gain. A generalized linear mixed model (gamma distribution, log link) confirmed that RTs varied systematically with reward magnitude, with participants responding fastest to high-reward trials and slowest to no-reward trials (*β* = -0.10; 95% CI [-0.11, -0.09], *χ*²(2) = 216.81, *p* <.0001; exp(β) = 0.90, 95% CI [0.89, 0.92], Fig. 4a), validating RT as a behavioral proxy for reward sensitivity. Additional effects of meeting (i.e. post vs. pre-training; *β* = -0.02; 95% CI [-0.03, -0.01]; exp(β) = 0.98, 95% CI [0.97, 0.99]; *χ*²(1) = 24.02, *p* <.0001), and run number (*β* = -0.03; 95% CI [-0.04, -0.01]; exp(β) = 0.98, 95% CI [0.96, 0.99]; *χ*²(1) = 12.24, *p* <.001) revealed that reaction times decreased with repetition both across runs and meetings.

**Figure 4.**
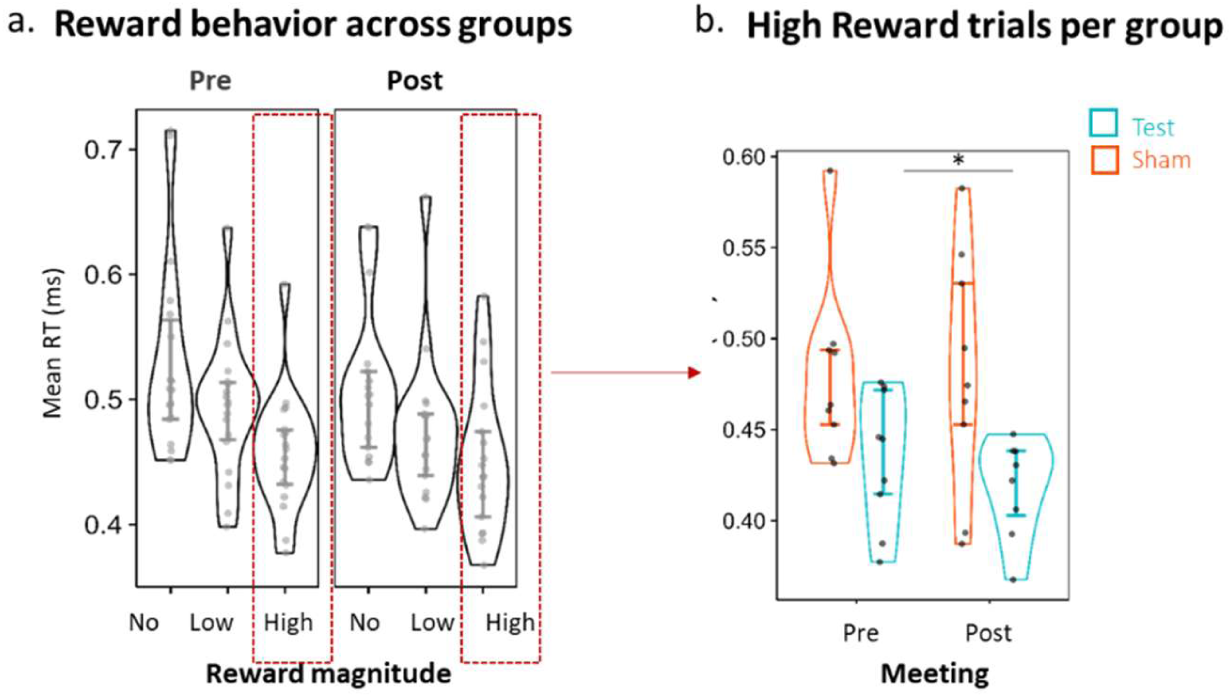
Study 2: functional training efficacy; response vigor in the CRRT (MID) task. (**a**) Mean Reaction times (RTs) as a function of anticipated reward magnitude, shown across groups (test, sham) and meeting (pre-, post-training), plotted separately for each reward level (no, low, and high reward). As expected, reaction times decreased with greater reward magnitude, confirming task sensitivity to reward. (**b**) Neurofeedback effects on reward behavior: Mean Reaction Times (RT) for high reward magnitude trials, plotted separately for each group and meeting. Individual data points represent mean RT values. Asterisks denote significant effects (*p* < .05). Error bars represent the standard error of the mean (SEM). Abbreviations: RT = Reaction Time, CRRT = Cued Reinforcement Reaction Task; MID = Monetary Incentive Delay

Critically, we tested whether neurofeedback training enhanced this reward-driven behavioral response during high-reward trials. A generalized linear mixed model (gamma distribution, log link) revealed a significant group-by-meeting interaction for RTs (*β* = -0.01; 95% CI [-0.02, -0.001]; *χ*²(1) = 5.28, *p* =.02; exp(*β*) = 0.99; 95%[0.978, 1.00]; Fig. 4b), indicating that reaction-time changes differed between groups. Specifically, the test group responded significantly faster after training compared to pre-training (post-hoc tests; post vs. baseline = -0.02, *z* = -3.07, *p* = .002), while the sham group did not show improvement (*z* = 0.07, *p* =.94). The model also revealed main effects of group (*β* = -0.06; 95% CI [-0.12, -0.01], χ²(1) = 4.93, p = .03), meeting (post vs. pre-training; *β* = -0.01; 95% CI [-0.02, -0.001], χ²(1) = 4.85, p = .03) and run number (*β* = -0.03; 95% CI [-0.06, -0.01], χ²(1) = 9.77, p = .002). These findings indicate differential training-related changes in high-reward reaction times between groups, with faster post-training responses observed in the test group only.

B.4. VS-EFP transfer learning is associated with improved control of the underlying VS-BOLD target The next key question was whether neurofeedback training for modulating the VS-EFP — an EEG-based signal model trained to reflect VS-related BOLD activation^42^ — relates to volitional control of the underlying fMRI BOLD target. To address this, we collected fMRI data before and after training while participants performed a self-neuromodulation task without feedback (transfer conditions). During scans, participants first rested for 120 seconds, then attempted to upregulate their brain activity for 120 seconds, similar to the transfer runs during EFP neurofeedback training. We quantified improvement in VS-BOLD regulation as the difference in beta-weighted right and left VS activity during the *regulate condition* after vs. before training. A mixed linear model analysis with group and hemisphere (right, left) as fixed effects and participants as random intercepts revealed that the test group showed greater improvement in VS-BOLD regulation compared to the sham group (main effect of group: *β* = 1.27; 95% CI [0.18, 2.36], *F*(1,12.71) = 5.69, *p* = 0.03; η² = 0.31). With no hemisphere effect (*β* = 0.13; 95% CI [-0.41, 0.67], *F*(1,13.08) = 0.25, *p* = 0.63), we averaged activity across bilateral VS for subsequent analyses (Fig. 5a). Descriptively, one-sample Wilcoxon signed-rank tests confirmed that only the test group significantly improved in upregulating VS-BOLD activation following training (*W* = 32, *p* = 0.03, one-sided), while the sham group did not (*W* = 17, *p* = 0.58, one-sided). Critically, improvement in VS-BOLD regulation correlated with VS-EFP transfer capacity during late-phase neurofeedback training (Fig. 5b). Specifically, in the test group, participants who showed greater VS-EFP transfer modulation during training also showed greater improvement in VS-BOLD regulation (*ρ* = 0.71, *p* = 0.03, one-sided), whereas no such relationship was observed in the sham group (*ρ* = 0.17, *p* = 0.35, one-sided). These findings suggest that VS-EFP neurofeedback training relates to volitional control of the underlying VS target, with late-phase VS-EFP learning correlating with fMRI-assessed improvement in VS self-neuromodulation.

**Figure 5:**
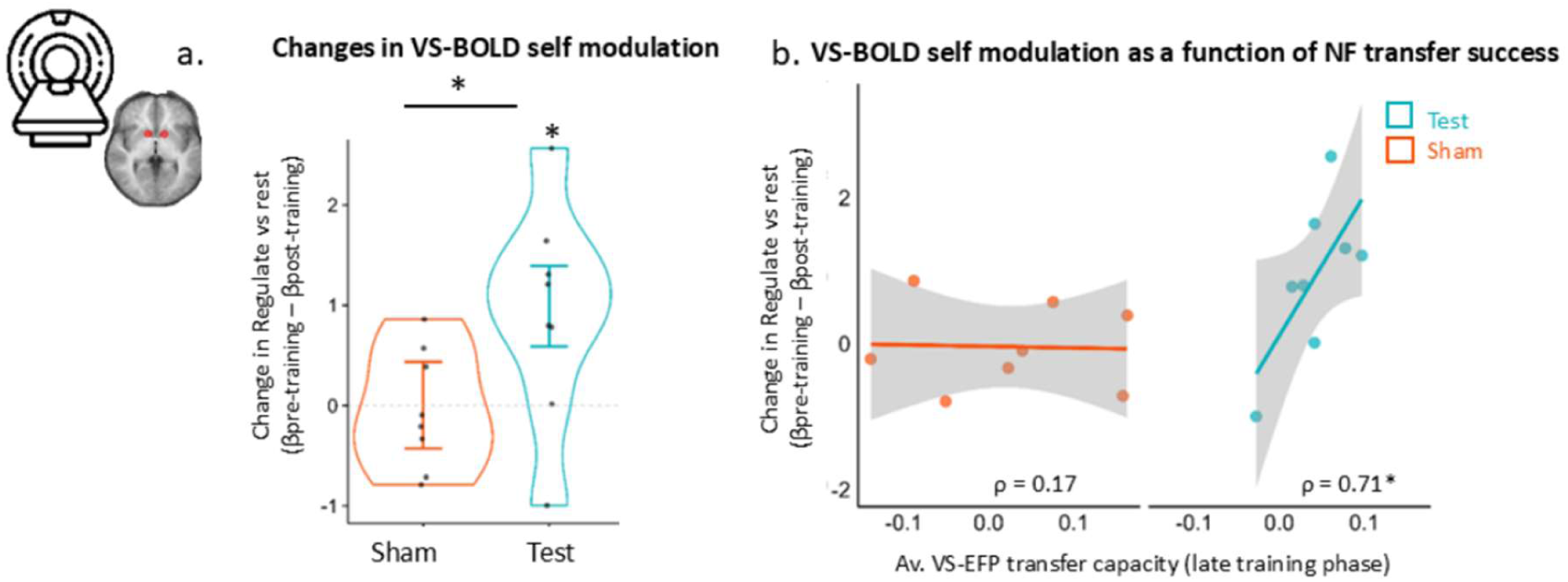
Pre-post changes in VS-BOLD self-regulation following neurofeedback training and their relationship to training performance. Change in VS-BOLD self-regulation following training for the test (cyan-blue, n=8) and the sham (orange, n=8) groups. VS-BOLD self-regulation ability was assessed using a pre-post fMRI design with a transfer task, during which participants were instructed to regulate their brain activity without external feedback. Change in left and right VS regulation was indexed as: [β regulate post-training] – [β regulate pre-training]. (a) Change VS-BOLD self-regulation ability per group shown as violin plots overlaid on individual data points; error bars represent standard error of the mean (SEM). (b) Correlation between VS-EFP transfer capacity during the late training phase and change in VS-BOLD self-regulation capacity, shown per group with regression lines and 95% confidence intervals. Asterisks denote significant effects; asterisks above violins indicating differences from zero (p < .05). Abbreviations: VS = ventral striatum; BOLD = blood-oxygen-level-dependent; EFP = Electrical FingerPrint;

### C. Pre-registered replication and mechanistic insights (Study 3)

We next designed a preregistered study (*N* = 40) to validate the Study 2 findings in a larger sample. We further optimized the protocol using the same neurofeedback setup and music manipulation as Study 2. First, to address the potential incidental contingency between VS-EFP and sham feedback observed in Study 2, we introduced a reversed sham condition in half of the sham participants (n = 10), whereby the yoked EFP signal was presented in reversed temporal order prior to feedback delivery. This maintained the proportion of positive versus negative feedback and distortion levels while reducing incidental temporal contingency arising from shared training dynamics. Second, to control for potential fatigue effects, the transfer cycle was placed first in each training session rather than last, allowing assessment of generalization from the previous training session without fatigue confounds. Third, to reduce repetition and maintain engagement, we increased the neurofeedback song pool from five to 10 songs, randomly selecting five per session (versus repeating the same five in Studies 1-2).

C.1. Contingent neurofeedback is associated with greater VS-EFP upregulation across training sessions. We first aimed to replicate the demonstration from study 2 that individuals can upregulate the VS-EFP better with contingent neurofeedback. Descriptively, one-sample Wilcoxon signed-rank tests on average neuromodulation capacity per training session confirmed that VS-EFP neurofeedback, but not sham, was associated with significant VS-EFP upregulation across all six training sessions (test group: *q*(FDR) < .002 for all sessions; sham group: *q*(FDR) >0.09; Fig. 6a).

**Figure 6:**
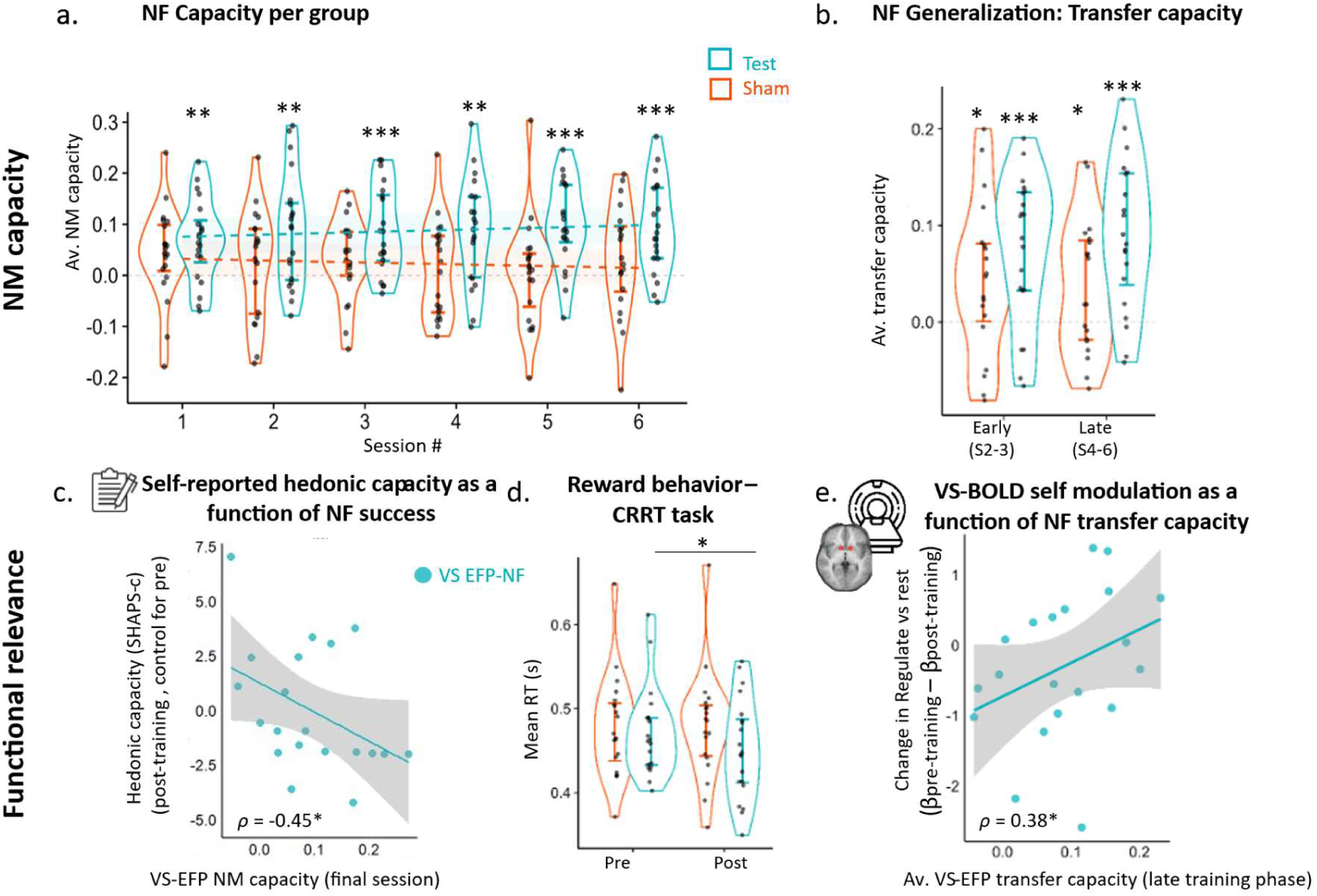
Study 3 - Pre-registered Validation Study in an Independent Data Set (n=40). **(a–b)** *Neurofeedback performance:* shown using violin plots with individual data points. **(a)** Neuromodulation (NM) capacity across cycles, shown per training session and separately for the test group (cyan-blue, n=20) and the control group (orange, n=20). Regression lines superimposed on the plots represent the linear trends (slope of learning across sessions) per group with corresponding colors. **(b)** *Generalization*: transfer capacity, shown per group and training phase. *Process-relevance***: (c)** correlation between neuromodulation capacity during the last session and post-training hedonic capacity, controlling for baseline hedonic capacity, plotted with regression lines and 95% confidence intervals. **(d)** Reaction times in the CRRT task for high-reward trials, plotted separately for each group and meeting (pre-training, post-training). *VS-EFP and VS-BOLD regulation relationship*: **(e)** Correlation between VS-EFP transfer capacity during the late training phase and change in VS-BOLD self-regulation activation during transfer trials (post versus pre-training). Asterisks denote significant effects (*p* < .05). Error bars represent standard error of the mean. Regression lines with 95% confidence intervals are shown for correlation plots. Abbreviations: NF = NeuroFeedback; NM = NeuroModulation; VS = ventral striatum, EFP = electrical fingerprint; SHAPS = Snaith-Hamilton Pleasure Scale. CRRT = Cued

Linear mixed-effects models revealed a significant main effect of group (*β* = 0.03, 95% CI [0.01, 0.06], *F(1, 18.99)* = 6.94, *p* = 0.02, η² = 0.27), indicating greater neuromodulation capacity in the test group versus sham. A significant group × session interaction with a linear trend was also observed (β *(linear)* = 0.02, 95% CI [0.001, 0.03], *F(5, 1118.99)* = 2.97, p = 0.01, η² = 0.013), reflecting increasing divergence between groups across training. Post-hoc comparisons showed that the test group outperformed the sham group beginning at training session 2 (*p* < .05). Post-hoc trend analyses revealed a significant positive linear increase in neuromodulation capacity across sessions in the test group (estimate = 0.16, SE = 0.08, *p* = 0.044), whereas no significant linear trend was observed in the sham group (estimate = −0.12, SE = 0.08, *p* = 0.12). These findings indicate that the test group showed greater VS-EFP upregulation than sham, with performance improving across training sessions.

To address the contingency concern from Study 2, Study 3 included two sham feedback variants (reversed vs. veridical). An analysis of each paired group separately confirmed that incidental temporal dynamics in the veridical sham condition may have blunted the group differences. Nevertheless, since a mixed-effects model revealed no difference in neuromodulation capacity between reversed and veridical (non-reversed) sham conditions (group effect: *β* = -0.02, CI [-0.09, 0.06], *F*(1, 17.85) = 0.19, *p* = 0.67), we pooled their data and did not distinguish between sham conditions in further analyses.

C.2. Neurofeedback is associated with greater overall transfer capacity. We next examined whether transfer capacity findings replicated when the transfer cycle was presented first in the training session. Descriptively, Wilcoxon signed-rank tests confirmed that VS-EFP neurofeedback was robustly associated with enhanced transfer capacity during late training (test group: *W* = 199; *p* < .001; Fig. 6b). Unlike Study 2, significant transfer capacity was also evident during early training in the test group (*W* = 192; *q*(FDR) < .001) and even in the sham group (early: *W* = 147; *q*(FDR) = .04; late: *W* = 143; *q*(FDR) = .04). Linear mixed models revealed that while the group-by-training phase interaction did not replicate (*β* = -0.01; 95% CI [-0.04, 0.01], *F*(1,147.92) = 1.22, *p* = 0.27), main effect of group emerged (*β* = 0.04; 95% CI [0.001, 0.04]; *F*(1, 15.9) = 6.23, *p* = .02, η² = 0.28). This suggests that VS-EFP neurofeedback, relative to sham, was associated with greater overall transfer capacity across training phases.

C.3. Greater VS-EFP upregulation is associated with post-training hedonic capacity and improved reward-driven behavior. We next validated the observed association between neuromodulation capacity and self-reported hedonic capacity, measured via the SHAPS questionnaire. Greater neuromodulation capacity during the final training session was associated with higher post-training hedonic capacity, corrected for baseline (*ρ* = -0.45, *p* = 0.05; Fig. 6c), replicating Study 2 and confirming the relevance of the neurofeedback training to self-reported hedonic capacity.

We next examined whether the behavioral effects observed in Study 2 replicated in Study 3. Pre-registered analyses of the PST and EEfRT, which showed training-related effects in Study 2, did not replicate in Study 3 (see Supplementary Information for full results). In contrast, analysis of the CRRT revealed a replicated effect, with faster reaction times in response to high reward. Although this analysis was not pre-registered, it is reported here for completeness. Specifically, a significant group-by-training session interaction emerged in reaction times during high-reward trials (*β* = - 0.01; 95% CI [-0.02, -0.001]; *χ*²(1) = 8.95, *p* = .003; exp(*β) =* 0.99; 95% CI [0.984, 0.997]).

Post-hoc analysis confirmed that the test group responded faster to targets following training relative to baseline (post vs. baseline = -0.02; *z* = -4.52, *p* < 0.001) while the sham group did not, (post vs. baseline = -0.002; *z* = -0.37, *p* =.71; Fig. 6d), replicating enhanced reward responsiveness following VS-EFP neurofeedback.

C.4. VS-EFP transfer learning is associated with improved control of the underlying VS-BOLD target. We next aimed to validate study 2 findings suggesting an association between VS-EFP and VS-BOLD regulation under transfer conditions. The results revealed a partial replication. While overall improvement in regulating VS-BOLD following training did not show the anticipated group effect (*β* = 0.23, CI 95% [-0.18 – 0.63], *F*(1, 31.25) = 1.3, *p* = 0.27), Spearman’s correlation confirmed that improvement in VS-BOLD upregulation following training was significantly associated with VS-EFP transfer upregulation during late training in the test group (*ρ* = 0.38, *p* = 0.05, one-sided; Fig. 6e), but not in the sham group (*ρ* = -0.22, *p* = 0.21). This replicates Study 2, supporting the link between EEG-based neuromodulation using the VS-EFP and target VS-BOLD activation.

C.5. Neurofeedback training is associated with enhanced auditory-striatal connectivity. Finally, in Study 3 we explored the mechanisms underlying musical VS-EFP neurofeedback. We hypothesized that learning to upregulate the VS-EFP signal with a musical interface would be associated with enhanced functional connectivity between the auditory cortex - the input region - and the VS^11,43^. We analyzed fMRI data from a music listening task conducted before and after training. Participants listened to two types of pleasurable, self-selected music: neurofeedback music (music used during training) and non-neurofeedback music (different music not used during training). We examined whether training enhanced connectivity between superior temporal gyrus (STG; auditory cortex) and VS during listening to trained vs. untrained music. For each participant, meeting (pre-training, post-training), and hemisphere (left, right), we calculated a music-selective connectivity index (ΔPPI = STG-VS connectivity during ‘neurofeedback music’ minus ‘non-neurofeedback music’). A mixed-effects ANCOVA tested whether post-training music-selective connectivity differed between groups, controlling for pre-training levels. The analysis revealed a Group-by-Hemisphere interaction (*β* = -0.08, *F*(1,30.9) = 5.58, *p* = 0.02, η² = 0.15). Post-hoc comparisons revealed that the test group exhibited significantly greater music-selective connectivity in the right STG-VS pathway compared to the sham group (test vs. sham = 0.41, *t*(46.5) = 2.73, *p* = 0.009), with no group difference in the left pathway (*p* = 0.63; Fig. 7). Descriptively, the test group showed positive right-hemisphere music-selective connectivity after training (Median = 0.39; Wilcoxon one-sample signed-rank test; *W* = 101; *p* = 0.04), indicating stronger connectivity during trained versus untrained music (Fig. 7), while the sham group did not (Median = -0.16; *p* = 0.99). These findings indicate that neurofeedback training was associated with enhanced right auditory-striatal connectivity specifically for music used during training.

**Figure 7.**
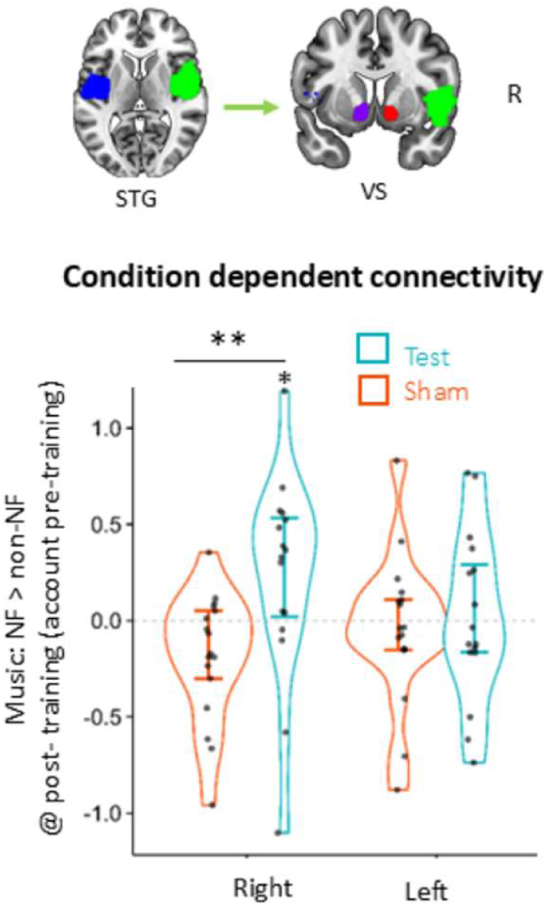
Study 3. Music-selective STG-VS functional connectivity following neurofeedback training. (**Top**) Seed and target regions overlaid on a canonical MRI brain (coronal view), plotted using CONN toolbox. Seed regions (right and left superior temporal gyrus, R. STG and L. STG) shown in green and blue; target regions (right and left ventral striatum, R. VS and L. VS) shown in red and purple. **(Bottom)** Music-selective connectivity between STG and VS following training, shown separately for test (cyan-blue, n=17) and sham (orange, n=15) groups. Left side: R. STG to R. VS connectivity; right side: L. STG to L. VS connectivity. Music-selective connectivity represents the difference in STG-VS connectivity during neurofeedback music versus non-neurofeedback music listening post-training, controlling for baseline. Violin plots overlaid with individual data points; error bars represent standard error of the mean (SEM). Asterisks denote significant group differences (p < .05); asterisks above violins indicate differences from zero. Abbreviations: NF = NeuroFeedback; VS = ventral striatum, STG = Superior Temporal Gyrus.

## Discussion

The current study demonstrates the feasibility and behavioral relevance of a novel musical neurofeedback approach for upregulating reward-related brain activity. Across three sham-controlled, double-blind studies, healthy individuals successfully learned to upregulate the VS-EFP signal through music-based neurofeedback (Studies 1-3); this neuromodulation capacity transferred to no-feedback contexts (Studies 2-3; Fig. 3c, 6b) and improved across repeated training sessions (Study 3; Fig. 6a). Greater neuromodulation capacity correlated with enhanced post-training self-reported hedonic capacity levels (Studies 2-3; Fig. 3d, 6c), and training resulted in faster reaction times to reward cues (Studies 2-3, Fig. 4b, 6d), indicating behavioral relevance. FMRI further revealed that VS-EFP neuromodulation capacity during transfer correlated with improved VS-BOLD upregulation under similar no-feedback conditions, validating this EEG-based marker as a scalable probe for reward-related self-regulation (Studies 2-3, Fig. 5b, 6e). Finally, neurofeedback training enhanced functional connectivity between right auditory cortex and VS during listening to trained music (Study 3, Fig. 7), suggesting that mechanistically, music-based neurofeedback may strengthen auditory-reward pathways that support VS engagement.

Feasibility and specificity of musical neurofeedback. Our findings demonstrated that musical neurofeedback targeting the VS-EFP was feasible, learnable, and specific, addressing key challenges in reward-focused neurofeedback. Prior real-time fMRI neurofeedback studies targeting reward regions^24–30^ showed feasibility but often reported limited transfer beyond training contexts^24,25,44^, or failed to demonstrate sustained modulation^24,27^. Our findings showed that repeated musical VS-EFP neurofeedback supported both learning and transfer, underscoring the value of multi-session approaches. Given the cost and limited scalability of fMRI-based neurofeedback used in prior reward self-regulation studies, our EEG-based approach offers a more accessible alternative for repeated training.

A novel element in our design was the use of pleasurable, self-selected music as the feedback interface. Unlike conventional neutral feedback signals (e.g., bar graphs) ^25–27,44^, this interface is both informative and affectively engaging, potentiating reward processes through a stimulus known to robustly activate the reward system^1,45,46,5,9,2^. This strategy aligns with process-based neurofeedback approaches advocating functionally relevant feedback^21^, which have been associated with enhanced learning and transfer^33,47^. We dynamically modulated the music’s acoustic properties to create a closed positive feedback loop, such that better self-regulation yielded more rewarding music. Personalization through self-selected music further tailored the experience to individual preferences, potentially enhancing reward responses^9,48^ and neural engagement^34,49^.In the present study, our priority was to maximize reward-related modulation through the use of a personally rewarding and modifiable stimulus; future work comparing musical and neutral feedback will help quantify music’s specific contribution.

In Study 2, although the test group significantly upregulated the VS-EFP signal, this neuromodulation capacity was not significantly greater than that of the sham group, which showed high variability. Post-hoc analyses revealed that signal-feedback contingency, even when incidental, predicted neuromodulation capacity in the sham group. This effect likely emerged from shared regulation dynamics between yoked pairs (e.g., onset responses, fatigue-related decay). After addressing this issue in Study 3 using a reversed sham condition, clear group differences emerged. Viewing neurofeedback as a reinforcement learning procedure^50,51^, these findings suggest that even partial contingency may influence self-regulation^52^, although more systematic examination of this phenomenon is warranted.

Behavioral relevance to reward processing. Across all three studies, VS-EFP neuromodulation capacity correlated with higher self-reported hedonic capacity (i.e., ability to experience pleasure), highlighting its clinical potential. VS-EFP training also resulted in faster reaction times to reward cues, consistent with a prior VS-targeted real-time-fMRI neurofeedback study^26^. Given that reaction time modulation by reward magnitude in such tasks reflects momentary reward sensitivity^26,53^, motivational engagement^54^, and VS activation^55^, the observed group specificity in the current study suggests that VS-EFP training enhanced reward sensitivity or motivation during reward anticipation. However, not all behavioral effects were replicated in Study 3 (effort exertion, probabilistic learning), underscoring the need for further replication and generalization to clinical populations.

Validation of VS-EFP as a target for reward-related self-regulation. In both Studies 2 and 3, VS-EFP neuromodulation capacity during transfer correlated with improved VS-BOLD modulation in the test group, confirming that EEG-based training engaged the original fMRI-defined target. Previously, we demonstrated that the VS-EFP, an fMRI-informed EEG model trained on simultaneous EEG-fMRI data during music listening, predicts VS-BOLD activation and is modulated by pleasurable music^42^. The current findings extend this work by showing that volitional VS-EFP neuromodulation correlates with VS-BOLD self-neuromodulation, strengthening the case that VS-EFP captures neural dynamics linked to VS engagement and can be utilized in neurofeedback. Converging evidence from neurofeedback for emotion regulation using an amygdala-related electrical fingerprint^56^, an fMRI-inspired EEG model, further supports this approach^57–62^.

Auditory-striatal connectivity as a mechanistic pathway. Growing evidence highlights the role of functional connectivity between auditory cortex and reward regions in musical pleasure^2,5,10,11,45,63–65^. Individuals with music-specific anhedonia exhibit weaker connectivity between these regions^10,66^, and moment-to-moment pleasure correlates with stronger auditory-reward coupling^2^. We hypothesized that this musical reward pathway could provide access to the reward system during neurofeedback. Study 3 revealed that, following training, the test group, but not the sham group, showed enhanced functional connectivity between right STG and right VS during listening to trained (i.e., songs used during neurofeedback) versus untrained music (Fig. 7). This right-lateralized effect is consistent with the right hemisphere’s predominant role in music processing^11^ and with evidence that right STG-VS functional connectivity is critical for music-induced pleasure^66^. This preferential enhancement for trained music emerged only in the test group, suggesting neurofeedback potentiated auditory-reward pathways through repeated pairing of music with successful VS modulation.

Recent work showed that receptive music interventions in older adults increase auditory-reward connectivity during music listening^67^, and that TMS-induced modulation of the striatum modifies auditory-reward connectivity and musical pleasure^43^. Extending this work, we found enhanced auditory-VS connectivity for music used in neurofeedback (vs. untrained music) during post-training listening without neurofeedback, indicating pathway strengthening beyond the training context.

These findings suggest that auditory-reward connectivity is modifiable through volitional training, extending the auditory-reward interaction model^11^ and revealing a potential mechanism through which music-based neurofeedback may enhance reward engagement.

### Clinical implications

Although conducted in healthy individuals, our behavioral measures—particularly hedonic capacity—are highly relevant to anhedonia, a core symptom of depression characterized by reduced pleasure in previously rewarding activities^23^. Given that anhedonia is consistently associated with diminished VS engagement^74^, interventions that directly modulate VS function are therapeutically relevant. A recent proof-of-concept study in patients with major depression and anhedonia using VS-EFP neurofeedback reported significant symptom reductions^69^, although it lacked a control condition. Our work demonstrated, with a sham control, that music-based VS-EFP neurofeedback produced learning and reward-specific effects.

Music-based neurofeedback may offer unique advantages, as general and musical anhedonia can dissociate in healthy individuals^70^, a pattern that could allow music to engage reward circuitry even in people with general anhedonia. Strengthening reward circuitry may also have implications beyond anhedonia, including stress buffering^71,72^, treating apathy in Parkinson’s Disease^22^, managing pain^73^, and addressing substance use disorders^22,24,28^. Complementary to neurofeedback, passive approaches using real-time EEG to personalize music selection have enhanced music-induced pleasure and chills^74^, suggesting multiple avenues for neurally guided musical interventions targeting reward function. Our results highlight the potential of repeated, reward-focused, music-based VS-EFP neurofeedback as a scalable intervention for anhedonia and reward dysregulation, warranting examination in clinical populations.

### Limitations

Despite methodological strengths, including double-blind, sham-controlled design and pre-registered replication, several limitations warrant consideration. First, not all outcomes replicated consistently; effects on EEfRT, PST, and group-level VS upregulation during transfer scans showed variability, possibly reflecting methodological differences or effect instability. Second, while the yoked sham design controlled for acoustic manipulation and subjective pleasure, it lacked feedback contingency, potentially reducing sham group motivation or inducing frustration^21,75^. Future studies should incorporate a contingent control condition targeting an alternative brain signal^21^, which would also allow more functional specificity. Third, without a neutral feedback interface, the specific contribution of pleasurable music remains unquantified. Additionally, ceiling effects in healthy individuals may have limited improvement, potentially underestimating clinical utility. Finally, outcomes were measured only immediately post-training; longer-term follow-up is needed to assess effect durability ^76,77^.

### Conclusions

Individuals can learn to self-regulate VS-related activity through music-based VS-EFP neurofeedback. Training effects were specific to the test group, transferred to no-feedback contexts, correlated with VS-BOLD, and were accompanied by enhanced auditory-reward connectivity. These findings offer mechanistic support for the auditory-reward interaction model^11^ and suggest that music-based neurofeedback acts by potentiating functional pathways linking perceptual and reward systems. This work establishes the feasibility and mechanistic validity of this process-relevant, scalable, personalized, reward-centered neurofeedback paradigm, laying the groundwork for applications in clinical populations with impaired reward function. Future work should include long-term follow-up and comparisons with alternative feedback interfaces to evaluate the durability, specificity, and generalizability of this intervention.

## Materials and Methods

### Study Design and Participants

NIH trial registration number (study 3): NCT04876170 (https://clinicaltrials.gov/study/NCT04876170)

### Study Design

We conducted three sham-controlled, double-blind neurofeedback studies with healthy volunteers. Study 1 (*N*=20; 10 test, 10 sham) was conducted at Tel-Aviv Sourasky University Medical Center and comprised a single neurofeedback session to assess feasibility of VS-EFP signal up-regulation using musical neurofeedback. Study 2 (*N*=20; 10 test, 10 sham) was conducted at the Montreal Neurological Institute, McGill University as a proof-of-concept longitudinal study consisting of six neurofeedback training sessions over 2-3 weeks (∼1-3 sessions per week), with pre- and post-training fMRI assessments to evaluate learning, generalization, and neurobehavioral outcomes. Study 3 (*N*=40; 20 test, 20 sham) was a preregistered replication study (NCT04876170; (https://clinicaltrials.gov/study/NCT04876170) conducted at Tel-Aviv Sourasky

University Medical Center using the same six-session training protocol as Study 2, with methodological refinements (see below). All studies compared real-time VS-EFP neurofeedback (test group) to yoked sham neurofeedback (sham group), where participants received feedback based on another participant’s VS-EFP signal (see Fig. 1 for study design overview).

#### Participants

In all three studies, healthy participants were recruited with no known history of psychiatric or neurological conditions (including hearing impairment). They were recruited through social media and across campus in all sites. All participants gave written informed consent before the beginning of the experiment and were compensated for their time. The experimental protocol has been approved by the Helsinki committee in Tel-Aviv Soursaky Medical Center (Studies 1 & 3; TLV-0401-17) and by the McGill Neurological Institute-Hospital Research Ethics Board (Study 2; MUHC, 2019-4769). <u>Study 1</u>: Twenty healthy Hebrew-speaking participants (11 females; mean age = 26.9 ± 3.99), who were randomly assigned either into test (N = 10; 6 females; mean age = 27.9 ± 4.59) or sham (N = 10; 5 females, mean age = 25.89 ± 3.11) groups. <u>Study 2</u>: Twenty healthy English-speaking participants (11 females; mean age = 21.1 ± 2.77 years), who were randomly assigned to test (N = 10; 6 females; mean age = 20.6 ± 2.12) or sham (N = 10; 5 females; mean age = 21.6 ± 3.47) groups. Two participants received both sham and real neurofeedback in different sessions due to technical difficulties. For these participants, only neurofeedback training data collected before group reassignment were included, and all post-training outcome data were excluded from analyses. Additional exclusions were applied on a per-analysis basis due to missing data, technical difficulties, or data quality concerns, including extreme outlier values (>3 interquartile ranges from the median in second-level analyses), a high rate of single-event rejection in nested datasets (greater than 1/3 of events), and excessive head motion in the fMRI data, defined as ≥20% of volumes flagged as motion outliers by the Artifact Detection Tools (ART) toolbox (https://www.nitrc.org/projects/artifact_detect/). The number of participants excluded per analysis was: neurofeedback training (neuromodulation: n = 0; transfer: n=1), SHAPS questionnaire (n = 2), fMRI transfer task (n = 4), EEfRT (n = 2), PST (n = 3), and CRRT (n = 2), resulting in final sample sizes of N = 20, 18, 16, 18, 17, and 18, respectively. <u>Study 3</u>: Forty-three participants were recruited. Three participants dropped out after the first pre-training baseline meeting, prior to randomization, resulting in a sample of forty healthy Hebrew-speaking participants (19 females; mean age = 25.03 ± 4.32 years), who were randomly assigned to test (N = 20; 10 females; mean age = 25.3 ± 4.62) or sham (N = 20; 9 females; mean age = 24.75 ± 3.97) groups. Exclusions per analysis, which followed the same criteria as Study 2, were neurofeedback training (n = 0), SHAPS questionnaire (n = 1), fMRI transfer task (n = 5), fMRI music task (n = 7), EEfRT (n = 7), PST (n = 0), and CRRT (n = 0), resulting in final sample sizes of N = 40, 39, 35, 33, 33, 40, and 40, respectively.

#### Sample Size Determination

Studies 1 and 2 were proof-of-concept investigations with modest sample sizes (*N*=20 each) designed to establish feasibility and identify methodological considerations. Study 3 sample size (*N*=40) was determined to double the proof-of-concept sample, providing adequate power for a preregistered replication (NCT04876170) while incorporating methodological refinements based on Study 2 findings (e.g., reversed sham feedback to minimize incidental contingency).

#### Randomization and Blinding

Participants were randomly assigned to either the test of sham groups at a 1: 1 ratio. Randomization took place following completion of the pre-training (baseline) meeting using a custom-made code implemented on the Open Vibe platform^78^. This code allowed for double-blinding between the test and sham groups by providing on-line feedback without revealing the source signal. Both participants and experimenters were blind to group allocation. Since the yoked sham method necessitates real data to be used for replay, full randomization started from the second participant in each cohort.

#### Group Allocation

Participants were randomly assigned to one of two conditions: (1) test (VS-EFP-neurofeedback; study 1: n=10; study2: n=10, study 3: n=20) (2) sham (yoked sham-neurofeedback; study 1: n=10; study2: n=10, study 3: n=20). The test group received continuous auditory feedback driven by their own VS-EFP amplitude changes. The sham group received continuous auditory feedback based on a sham-yoked method, wherein each participant from the sham group was paired to a participant from the test group, thus receiving the musical feedback of the paired test participant. This way, both groups were exposed to the exact proportion of sound manipulation that indicates their success-level, but only for the first group was it temporally related (i.e. contingent) to VS-related activity. In Study 3, half of the sham group received reversed yoked feedback to further reduce incidental contingency between the sham feedback and participants’ actual VS-EFP signals while maintaining the same levels of acoustic distortion and success rates.

### Neurofeedback Training Procedure

Musical stimuli selection and preparation. Participants were asked to provide a list of 10 to 15 of their favorite songs before coming into the lab for the experiment. They also indicated which section of the self-selected music they found most enjoyable (between 90 and 120 seconds in duration). Electronic digital versions of the song clips were cut down to 90-120 seconds around the indicated peak and edited in Audacity® (Audacity Team, version 2.0.5) to balance the sound level and remove any periods of silence prior to the beginning of the music. Five of the selected songs-in studies 1 + 2 and 10 of the selected songs from the list of favorite songs were used as the feedback in the neurofeedback training, with each of five different songs presented in each session, in random order. Five of the excerpts that were used during the neurofeedback training session (hereby termed neurofeedback songs), and five that were not used during neurofeedback (hereby termed non-NF songs), were further used in the fMRI music listening task using 50 sec excerpts.

VS-Related Electrical Fingerprint Model (VS-EFP). For the neural probe, we used an EEG model of reward-related activation related to the VS, termed VS-EFP, which was recently developed by our group and extensively validated^42^. This probe was developed by applying machine learning algorithms to predict the BOLD activation within the VS and related activation using the concurrently acquired EEG data. The modeling procedure resulted in a one-class model comprised of a time delay × frequency × weight x channels coefficient matrix. To extract the EFP, newly acquired EEG data recorded from eight electrodes (C4, F7, F8, T7, T8, P8, TP9 and TP10) at a given time point are multiplied by the coefficient matrix to produce the predicted VS-related fMRI-BOLD activity. The reliability of this generic model was validated in a series of analyses, revealing that the VS-EFP predicted BOLD activation within the VS and associated network in an independent sample, as well as under a different reward context^42^.

EEG data acquisition and online processing. In all studies, EEG data were recorded using a battery operated BrainAmp EEG amplifier (Brain Products, Munich, Germany) and a customized BrainCap electrode cap with Ag/AgCl ring electrodes providing 13 EEG channels and 1 electrocardiogram (ECG) channel, connected to an electrode input box (EIB-64 DUO, Brain Products, Munich, Germany). The electrodes were positioned according to the 10/20 system and included the eight electrodes of the VS-EFP model and additional channels (fp1, fp2, Fz, Cz, Pz), all in reference to the reference electrode between Fz and Cz. Raw EEG was sampled at 500 Hz.

#### Online calculation of VS-EFP

To provide continuous feedback based on momentary VS-EFP changes, we used an in-house neurofeedback platform built on OpenVIBE, an open-source brain-computer interface^78^. EEG signals were recorded via the OpenVIBE acquisition server and streamed in real time to a customized OpenVIBE Designer implementing the VS-EFP model. Real-time preprocessing included high-pass filtering (0.005 Hz) to remove slow drifts and notch filtering (50 Hz) to eliminate line noise.

The model operated on a continuously updated 30-second window of EEG activity, segmented into 1-second overlapping epochs (50% overlap). For each epoch, spectral features were extracted from predefined frequency bands and channels^42^. Features were normalized using mean and standard deviation parameters from a global baseline recording, ensuring real-time estimates reflected deviations from resting-state activity. The feature time series was low-pass filtered, downsampled by a factor of two, and projected onto trained model weights to compute VS-EFP amplitude. This computation updated every 3 seconds, enabling near real-time tracking of VS-related activity for continuous auditory neurofeedback. This neurofeedback platform was developed in-house based on our previous work and following similar principles^58^.

#### Musical Neurofeedback Interface

The neurofeedback interface used pleasurable, self-selected music to provide reward-related feedback. Study 1 employed simple amplitude manipulation. Studies 2 and 3 used a newly developed acoustic manipulation scheme to modulate musical hedonic value in a process-specific manner. This approach was developed and validated in pilot experiments to ensure real-time implementation on any sound file with multiple distinguishable reward value levels.

Sound quality was manipulated via band-pass filtering implemented in Max MSP (Cycling ‘74, Covina, CA, USA), updated every 3 seconds. A 512-band convolution FFT-based filter modulated bandwidth proportionally to VS-EFP neuromodulation success. Maximum quality (all bands active, full spectrum) corresponded to greatest pleasure; minimum quality (limited bands active, narrow spectrum) produced muffled sound resembling low-quality telephone audio.

In both cases, the self-selected music’s sound *volume* (study 1) or *quality* (studies 2+3) was modulated in real-time, every 3 seconds, and in monotonic correspondence to the EFP neuromodulation index, which quantified the deviation of the current VS-EFP value from the participant’s local baseline (the ‘listen’ block preceding self-neuromodulation where the participant was exposed to the feedback yet was asked not to perform any brain modulation). This neuromodulation index was calculated by subtracting the mean of the EFP values during the local baseline from the current EFP value and dividing the result by the standard deviation of the EFP values in the local baseline. This was achieved by applying the following equation:

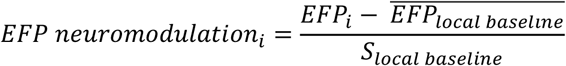

Where *EFP*_i_ is the current EFP value, representing the prediction of the targeted VS-related activation, 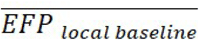 is the mean of the EFP values during the local baseline and the *S*_local baseline_ is the standard deviation of the EFP values during the local baseline. The increase or decrease in the music volume level or sound quality were in a monotonic correspondence with a *feedback value*; a rescaled value that of this EFP modulation index, which ranged from 0 to 1, where 0 represents the lower possible success value and 1 the highest possible value. To avoid saturation of the feedback with increasing success, the minimum and maximum success values used for calculating this rescaled feedback value were adapted from cycle to cycle based on the participant’s neuromodulation success in the previous cycle (initially restricted to a range between -2 and 2 and further adapted if exceeded). For the volume manipulation, the minimal value was set to 10% of the individual sound volume and the maximal value, to 90% of the individual sound volume. During the local baseline phase, participants heard their selected music softly (study 1, 5% of the maximal individual sound volume) or with low quality (study 2+3, i.e., highly filtered, 1% of the maximal value) without being able to influence its loudness or quality, respectively. In Studies 1 and 2, the local baseline consisted of passive music listening throughout, whereas in Study 3 music began only after 30 s of silence, matching the regulation phase in length. During the regulation phase, participants were instructed to try to make the music louder (study 1) or make it sound better (studies 2+3). To provide a smoother experience, the feedback value in studies 2+3 was smoothed over two time points using a moving average. This approach enabled participants to receive continuous, individualized neurofeedback reflecting ongoing changes in VS-related neural activity relative to their own baseline.

Neurofeedback Training Procedure. The training procedure is depicted schematically in Fig. 1. In study 1, the neurofeedback training included one session. In studies 2 and 3 the repeated neurofeedback training included six sessions, with a gap of two to four days between consecutive sessions. Each session lasted approximately 60 minutes, including preparation time, and began with a three-minute long ‘resting-state’ recording with eyes closed, which was henceforth used as a global baseline. This baseline data was used to compute normalization parameters and model-specific preprocessing coefficients, enabling subsequent real-time standardization and comparison of neural activity relative to each participant’s individual baseline. Following this baseline recording, participants were trained to upregulate their VS-EFP by receiving feedback on their success via the musical interface. The training paradigm across all sessions followed a similar design, composed of five repeated training cycles with eyes closed, each including two consecutive conditions: (a) *listen* (study 1: 150 sec.; Study 2 + 3: 120 sec) and (b) *regulate* (study 1: 120 sec.; Studies 2 & 3: 90 sec). During *listen,* participants were instructed to passively listen to the music, not to think of anything in particular and were informed that the sound they heard was not influenced by their brain activity. During *regulate,* participants were instructed to find the mental strategy that would make the music louder (study 1) or sound better (studies 2+3, see details below). In study 1, instructions were intentionally nonspecific, allowing individuals to adopt the mental strategy that they subjectively found most efficient. In Studies 2 and 3, to facilitate learning, instructions included a set of possible mental strategies that participants could explore in addition to their own. To keep the targeted process concealed, the proposed strategies involved mental activities related to music. Some of these were more connected to reward processing, such as recalling positive memories, practicing self-affirmation and motivation^27^ or imagining oneself immersed at a concert, while others were unrelated, like using motor and visual imagery of the musical performance. After each cycle, participants received a debriefing graph showing their regulation success, and underwent a series of computerized questions about the mental strategies they had used. During training, participants listened to the neurofeedback music at their preferred volume through headphones (Study 1: Sennheiser HD 202) or insert earphones (Studies 2 and 3: IP30 50 Ohm, RadioEar, New Eagle, PA). Prior to training, the volume was adjusted from a baseline of 50% (computer volume setting) to each participant’s preference during a brief sound-check run.

### Outcome Measures and Assessments

In all three studies, participants completed questionnaires assessing hedonic capacity, musical aptitude, and demographics before group assignment. Mood state was monitored before and after all training sessions.

In Studies 2 and 3, participants further underwent detailed pre- and post-training assessment meetings comprising additional questionnaires, fMRI scanning, and computerized behavioral tasks. fMRI protocols included structural scans, a neurofeedback transfer run (self-regulation without feedback), a music-listening task, and additional functional scans. Behavioral assessments included reward-based decision-making tasks. Physiological recordings (skin conductance in Study 2; EEG in Study 3) were also acquired during scanning but are outside the scope of this paper. Full details of all measures and protocols are provided below.

#### Self-Reported Questionnaires

Hedonic capacity was assessed using the Snaith-Hamilton Pleasure Scale^35^ (SHAPS; Study 2, English version: Cronbach’s α = 0.803; Study 3, Hebrew translated version: Cronbach’s α = 0.824). To capture inter-individual variability in hedonic capacity, continuous SHAPS scores were used (rather than dichotomous clinical cutoffs)^79^ by summing responses across all four response categories. Specifically, the 14 items were scored on a 4-point Likert scale (0 = strongly agree; 3 = strongly disagree), with higher total scores indicating lower hedonic capacity (i.e., greater anhedonic tendencies). Self-reports of mood (PANAS^80^), reward-related traits (BIS/BAS^81^, TEPS^82^, SPSRQ^83^, BMRQ^70^) and of musical training (MMHQ^84^) were also collected, using English (study 2) and Hebrew translated versions (study 3), yet they are outside the scope of this paper.

### Pre-Post Training Behavioral Tasks

Cued Reinforcement Reaction Time Task. Participants completed a modified version of the cued choice-reinforcement reaction time (CRRT) task^39,40^ during fMRI scanning, before and after neurofeedback training, to assess changes in reward responsivity. This task was selected because it provides a behavioral measure of reward vigor while neurally dissociating anticipatory and consummatory phases of reward processing. The key difference from the widely used Monetary Incentive Delay task^86^ is that in the CRRT, reward magnitude varies with reaction time performance through a dynamic algorithm, providing a more sensitive behavioral measure of reward responsivity^40,87^.

In each trial, participants were presented with a cue (750 ms) indicating the maximum possible monetary reward (Study 2: 0, 0.6, or 3 Canadian dollars; Study 3: 0, 1.2, or 6 Israeli Shekels). After a delay (2.5-3 s), participants rapidly identified an outlier among three circles by pressing the corresponding button. Feedback was displayed for 1.5 s, showing the amount won based on their reaction time. Following practice runs (12 trials outside scanner, 6 inside), participants completed two main runs of 32 trials each (7.5 min per run), one preceded by 30 s of silence and the other by a 30 s excerpt of neurofeedback music (order counterbalanced across participants). The CRRT was designed to capture both behavioral and neural indices of reward processing. However, the present paper focuses exclusively on behavioral reaction time data; fMRI data from this task are beyond the scope of this report.

#### Additional behavioral tasks

We also administered the Effort Expenditure for Rewards Task (EEfRT)^38^ to assess motivation for reward and the Probabilistic Selection Task (PST)^37,85^ to assess reward-based learning. Task details and results are reported in Supplementary Methods and Supplementary Results.

### Pre-Post fMRI Tasks

#### Transfer

To test for target engagement in the VS, and learning generalization (transferability) between contexts, participants in studies 2 and 3 underwent a transfer regulation task before and after training, while being scanned via fMRI. The transfer regulation task was the same as the one applied during the training sessions and was composed of 120 seconds of rest, followed by 120 seconds of regulation, with no feedback. Participants received the same instructions they got during the neurofeedback sessions, to rest and not think of anything in particular during the rest phase, and to apply a mental strategy during the regulation phase. During the pre-training meeting, participants received a list of proposed strategies (see description in the neurofeedback session) and were requested to apply one of them during the regulation phase. During the post-training meeting, participants were asked to choose the strategy that worked best during training.

#### Music Listening Task

To further test the musical neurofeedback learning mechanism, participants were engaged in a pleasurable music listening task before and after training, while undergoing the fMRI scan, which followed a similar design as applied in our previous studies^10,43^. During scanning, participants listened to 10 excerpts drawn from their favorite self-selected songs, with each excerpt from the part of the song they most enjoyed (each 50 seconds long). While listening, participants continuously rated on a scale from 1 to 4 (1 = neutral, 2 = low pleasure, 3= high pleasure, 4 = peak pleasure) how much pleasure they were experiencing at each moment using an MR compatible button box. The song presentation was interleaved with epochs of silence (each about 25 seconds, with a jitter of 3.5 seconds). The entire experimental paradigm lasted 12 minutes and 51 seconds. During listening, they were instructed to press a button corresponding to their rating for as long as they were experiencing that degree of pleasure. Importantly, to assess the specific effect of neurofeedback exposure, five of the songs were from a pool of songs that were used for the neurofeedback training, and the remaining five were only presented during scanning. We analyzed the data from study 3, as this was a mechanistic investigation following the initial replication. During the task, the audio files were played on a desktop computer using an in-house script utilizing Psychtoolbox and MATLAB software. Participants listened to the music through MRI-compatible insert earphones (earetrics Model S14, Sensimetrics Corporation, Malden, MA, USA) adjusted to a comfortable level while lying in the scanner.

Participants also underwent a resting state scan during which they asked to lie still and close their eyes while clearing their mind, and were also scanned while performing the CRRT task, but both of these datasets are beyond the scope of this paper.

#### fMRI data acquisition

Structural and functional scans were performed in a 3.0 Tesla Siemens MRI system (MAGNETOM Prisma, Germany) using a 32-channel (study 2) or a 20-channel (study 3) head coil. High-resolution T1-weighted anatomical images were acquired using a T1-weighted 3D Sagittal MPRAGE pulse sequence (study 2: TR/TE = 2300/4.18 ms, flip angle = 9°, voxel size = 1X1X1mm, FOV = 256 mm; study 3: TR/TE = 1860/2.74 ms, flip angle = 8°, voxel size = 1X1X1mm, FOV = 256 mm) was used. Functional whole-brain images were acquired in an interleaved top-to-bottom order, using a T2*-weighted gradient echo planar imaging pulse sequence (Study 2: TR/TE=780/33 ms, flip angle=52°, voxel size=2 X 2 X 2 mm, FOV=208 mm, 72 slices per volume; Study 3: TR/TE=3000/30 ms, flip angle=90°, voxel size=2.5 X 2.5 X 3 mm, FOV=220 mm, 43 slices per volume). In study 2, multi-band acceleration was applied with a factor of six, allowing simultaneous acquisition of multiple slices, thereby reducing the TR. The in-plane acceleration (GRAPPA) factor was two. Furthermore, to correct for distortions due to magnetic field inhomogeneities, field maps were acquired using a dual-echo gradient-echo sequence (TR/TE1/TE2 = 421/3.97/6.43 ms, voxel size = 3 X 3 X 3 mm³, matrix size = 74 x 74, field of view = 222 mm). The phase difference image was used to compute the B0 field inhomogeneity map, which was subsequently applied during preprocessing to correct for susceptibility-induced distortions in the functional images.

### Data Analysis

Offline neurofeedback Data analysis.

#### Preprocessing

EFP data exceeding a value of +-5 or 2.5 standard deviations from the mean of the entire signal were discarded. Cycles in which more than 20% of the data were considered noisy and discarded from further analysis.

NF performance indices and statistical analysis. Neurofeedback neuromodulation *capacity* (i.e., upregulation success) in each cycle and training session was measured as the mean difference in the signal power (VS-EFP) during the ‘regulate’ relative to the ‘listen’ (local baseline) condition^36,57^. The neurofeedback neuromodulation capacity of participant per cycle and session was analyzed using a linear mixed effects model with group (test versus sham), session number (for studies 2+3; 1-6, ordinal scale) and the group by session interaction as fixed effects. Cycle number was also added as a covariate to account for potential trends within session. Random intercepts were included for yoked-pairing and for individual subjects, nested within each yoked pair to account for the hierarchical structure of the data. Neurofeedback *Generalization capacity* was assessed as the neuromodulation capacity (VS-EFP signal difference: regulate - rest) during the *transfer trials*, separately assessed across the early- and late phases of the training sessions (sessions 2-3, and 4-6, respectively; early phase did not include session 1 since the transfer cycle was applied prior to training in session 1). The effect of training and phase was assessed using a similar linear mixed effects model, now with group, training phase, and their interaction as fixed effects, along with training session number, and the same definition of random intercepts. For study 1, a similar analysis, only without the session numbers, was applied. *Learning contingency,* the average correlation between participants’ brain activity (VS-EFP signal), and their received feedback during each session was also calculated for study 2. This measure was used in a linear mixed model for the control group only and in the linear mixed effects models described above, with the exception that instead of dichotomous group assignment, the continuous measure of learning contingency was used.

Outlier trials (cycles), below and above 1.5 * the Interquartile range of neuromodulation capacity across each group, were excluded prior to model fitting or averaging over sessions. In cases where the model with both intercepts for individual participants and the yoked pairs did not converge, we applied model selection based on AIC and BIC comparing two simpler models: one with random intercepts for pairs and second with random intercepts for individual participants, and selected the simpler model. There was one case in which the model with both intercepts for individual participants and the yoked pairs did not converge: the learning generalization analysis in study 2. The selected model in this case, with random intercepts for individual participants only, was preferred for its better balance of fit and simplicity.

### fMRI Data Analysis

#### Preprocessing

Functional and structural MRI data were preprocessed using SPM12 (Wellcome Centre for Human Neuroimaging, UCL, London, UK^88^). Preprocessing included slice-timing correction (multi-band acquisition in Study 2; interleaved in Study 3), realignment to correct for head motion, coregistration of structural to functional images, and spatial normalization to MNI152 space using deformation fields from tissue segmentation. Functional images were smoothed with a 6 mm FWHM Gaussian kernel and high-pass filtered [Transfer task: 0.0078 Hz]. Motion parameters were included as nuisance regressors. Field map-based distortion correction was applied in Study 2, in which such maps were acquired. For connectivity analyses, additional preprocessing and denoising steps were performed using the CONN toolbox (version 22.a, RRID:SCR_009550)^89^, including outlier detection (ART)^90^, motion/outlier regression, and high-pass filtering (>0.008 Hz).

#### Transfer task analysis

At the first level, a single general linear models (GLMs) across both pre- and post-training meetings was calculated per participant. The model included two task-related regressors per meeting: regulation prompt (brief cue to begin regulation) and regulation phase. The contrast of interest was regulation phase versus rest (local baseline). Given our a priori hypotheses regarding VS-BOLD engagement, we conducted region-of-interest (ROI) analyses. Bilateral VS ROIs were identical to those used to develop the VS-EFP model^42^, derived from a Neurosynth^91^ meta-analysis of the term “reward”. Average β coefficients within each ROI were extracted per participant and meeting using MarsBaR^92^. To quantify training effects, we calculated improvement in VS-BOLD regulation as the difference in VS activity during regulation vs. baseline post-training compared to pre-training [ΔVS-BOLD = β(post-training) - β(pre-training)].

Music listening task and connectivity analysis. A single GLM across both pre- and post-training meetings per participant included eight task-related regressors: neurofeedback music and non-neurofeedback music conditions (each with separate regressors for sustained listening [5 seconds after onset] and onset [first 5 seconds]), pleasure rating events with first-order parametric modulation reflecting rating magnitude, and motion parameters. The contrast of interest was sustained neurofeedback-music vs. non-neurofeedback music listening.

Functional connectivity was assessed using generalized psychophysiological interaction (gPPI^93,94^) analysis in the CONN toolbox. ROIs included bilateral VS (as above) and bilateral superior temporal gyrus (STG; auditory cortex), extracted from a meta-analysis of music-induced pleasure^5^. The ROIs are depicted in Fig. 7. For each seed-target pair, gPPI models included seed BOLD signal as the physiological factor, task condition regressors (convolved with canonical HRF) as psychological factors, and their product as psychophysiological interaction terms. Analyses focused on auditory cortex seeds and VS targets. Average β coefficients of PPI terms within each ROI were extracted per participant, meeting (pre-, post-training), and condition (neurofeedback music, non-neurofeedback music).

To quantify training-related changes in connectivity, we computed within-subject music-selective connectivity index per meeting: ΔPPI = β(neurofeedback music) - β(non-neurofeedback music), representing the difference in STG-VS connectivity during neurofeedback versus non-neurofeedback music. To assess training-related effects on music-selective connectivity, a mixed-effects ANCOVA was fit for predicting the post-training neurofeedback - non-neurofeedback connectivity contrast from Group (test, sham), side (left, right), and their interaction, while controlling for the corresponding pre-training contrast. Participants were included as a random intercept.

### Behavioral Data Analysis

Behavioral measures (CRRT, EEfRT, PST tasks), which were coded per trial, subject and meeting were each assessed with separate generalized linear mixed effects models with group (test, sham) and meeting (pre-training versus post-training) as fixed factors and participants as random intercepts. In each model, additional fixed factors of no interest, accounting for the variance in stimulus presentation or trial time were added. Trials with short reaction times (250 ms), or with incorrect or invalid responses, were identified and removed prior to the regression analysis. Individuals with rates of over 33% of invalid trials were excluded from the analysis. Behavioral results from the Cued Reinforcement Reaction Task (CRRT; reaction times) are reported in the main text, whereas results from the EEfRT (choice) and the Probabilistic Selection Task (PST; accuracy) are reported in the Supplemenary materials, where full analysis details are provided.

Correlation between performance and outcome measures. Accounting for baseline performance. To adjust for individual differences in baseline performance when evaluating the correlation with post-training effects, we accounted for the pre-training performance within each group. To achieve this, we used a linear model (lm function in R), where post-training performance was modeled as a function of pre-training performance and group. The residuals from this model were extracted and used for correlation analyses with the neurofeedback success measures. To not rely on normality assumptions, we employed Spearman’s rank correlation to examine these relationships.

In the ROI and correlation analyses, outlier data points exceeding 3 * interquartile range from the 25th or 75th percentiles were excluded from.

### Statistical Analysis

All statistical analyses were conducted using R (R version 4.3.3^95^) and SPM12^88^. Significance threshold was set at α = 0.05. For analyses involving multiple comparisons across sessions or cycles, false discovery rate (FDR)^96^ correction was applied using the Benjamini-Hochberg procedure, with corrected p-values denoted as q(FDR). All statistical tests were two-tailed unless otherwise specified (one-tailed tests were used for directional a priori hypotheses).

Linear mixed-effects models were estimated using the lmer function from the lme4 package^97^. Reaction times in the CRRT were modeled using a Gamma distribution and a log link function using the glmer function from the lme4 package.

In all models, fixed effects were evaluated using omnibus tests appropriate to the model class. For linear mixed-effects models, fixed effects were assessed using Type III analysis of variance with Satterthwaite’s approximation for degrees of freedom, whereas for generalized linear mixed-effects models, fixed effects were evaluated using Wald χ² tests. Type III tests were used to evaluate main effects and interactions in models including interaction terms. In the absence of interaction, Type II tests were used. In addition to omnibus tests, model coefficients were extracted from model summaries, and effect sizes are reported as regression coefficients or odds ratios with 95% confidence intervals. When models failed to converge with the full random effects structure, model selection was performed using Akaike Information Criterion (AIC) and Bayesian Information Criterion (BIC).

Significant interactions were decomposed using planned post-hoc contrasts based on estimated marginal means (emmeans package^98^), with Holm correction for multiple comparisons.

Non-parametric tests were used for supplementary or descriptive analyses to complement the primary inferential models and are reported separately. For these tests, we report test statistics (W) and p-values.

## Supporting information

Task details and results are reported in Supplementary Methods and Supplementary Results file

## Acknowledgments

We would like to thank all participants for their time. We also thank Nathaniel Lynch, Lyla Hawari, Liam Pantis, Isabel Levine, Noam Sarna, Shir Caspi, Shir Shintel and Diana Arbich for their assistance with data collection and pilot testing. We also thank Yashar Zeighami and Matthias Kirschner for their advice and guidance regarding the computerized reward tasks as well as for Ernest Mas-Herero for his advice on the music pleasure task. We also thank Shimon Shahar and Aya Vitury for valuable advice on the linear mixed modeling approach.

