## Supplementary material for "Music-to My Brain: Modulation of Reward System Activity via Musical Neurofeedback": Task details and results are reported in Supplementary Methods and Supplementary Results file

---

Table of Contents

Supplementary Methods........................................................p.2

Supplementary Results *(including Figures S1-S4 and Tables S1-S4)*........................................p.8

#### Supplementary Methods

*Computerized Behavioral Tasks*

Cued Choice-Reinforcement Reaction Time (CRRT). *Task overview and procedure*: Participants underwent fMRI scanning while performing a modified version of the *CRRT* task[1,2]. This paradigm was designed to asses, in additional to the neural signatures of the response to reward anticipation and consumption, behavioral indices of reward responsivity and vigor in anticipation of reward, as reflected in reaction times (RTs) that scale with reward magnitude[2,3]. The CRRT follows the general structure of the *Monetary Incentive Delay (MID)* task[4], with the key difference that the actual reward is contingent on each participant's reaction time rather than on trial accuracy alone.

Each trial began with the presentation of one of three geometric cues (750 msec), indicating the maximum potential monetary reward in that trial, being high, low or none (Study 2: 0.6 ¢, 3 CAD, 0 ¢; Study 3: 1.2 ILS, 6 ILS, 0 ILS). After a variable delay (2.5-3 s), participants identified an outlier among three circles displayed to the left or right of fixation by pressing the corresponding button. Immediately following the response, feedback was presented for 1.5 s, showing the amount of money won on that trial, displayed next to a red horizontal line.

The reward magnitude in the CRRT was determined by a *dynamic algorithm* that continuously adjusted the monetary feedback according to each participant's recent performance, thereby providing a sensitive measure of reward-driven motor vigor. Participants completed one practice run outside the scanner (12 trials) and one short practice run inside the scanner (6 trials) prior to the main experiment; neither was included in the data analysis. The task consisted of two runs of 32 trials each (≈7.5 min per run). One run was preceded by 30 sec of silence and the other by 30 sec of the musical excerpt used during neurofeedback (counterbalanced across participants) to reinstate the training-related state. The present analyses focus exclusively on behavioral RT measures.

Statistical analysis: To examine how reaction times (RTs) reflected reward responsivity and training-related changes in motor vigor, we fitted two generalized linear mixed-effects models (GLMMs) using the the glmer function from the lme4 package in R[5]. Because RTs were positively skewed and non-normally distributed, they were modeled with a gamma distribution and an log link function. All models included a random intercept for subject to account for repeated measures. To confirm that RTs varied systematically as a function of reward magnitude, we fitted the a model, where RewardLevel (high, low, none) was modeled as a fixed ordered effect of interest, and Meeting Number (post/pre), Music Played Before (yes/no), Trial Number, and Answer Side (right/left) were included as covariates of no direct interest.

To assess whether neurofeedback training modulated reward-driven response vigor, a second model was fitted to high-reward trials only where we modeled the interaction between group assignment (Test or sham) and session (pre- vs. Post-training) as fixed effects of interest, while Meeting Number, Music Played Before, Trial Number, and Answer Side were included as covariates of no direct interest. Model performance and fixed-effect significance were assessed using Wald $\chi^2$ tests, and model fit was summarized with marginal and conditional $R^2$ values.

Effort-Expenditure for Rewards Task (EEfRT). *Task overview and procedure:* This task was used to evaluate incentive motivation behavior by examining participants' willingness to choose higher-effort, higher-reward options rather than lower-effort, lower-reward alternatives[6]. This is a multi-trial task in which participants are asked to choose between an easy or a hard task for the chance of gaining a monetary reward of a given magnitude and gain probability. In each trial, participants are asked to choose between a hard-task and an easy-task. Completion of the task may lead to a monetary reward. Successful completion of the hard task required the participant to make 100 button presses, using the non-dominant little finger within 21 seconds. If they succeed, participants were eligible to win an amount of money within a range of 1.2 to 4 Canadian dollars (CAD; study 2) or 2.5-8.5 Israeli Shekels (ILS; study 3). Successful completion of easy-task trials required the participant to make 30 button presses, using the dominant index finger within 7 seconds. For this choice of the easy condition, participants were eligible to win 1 CAD or 2 ILS if they successfully completed the task. Participants were not guaranteed to win a reward if they completed the task; they were provided at the beginning of each trial with the probability of gaining the money if the task was completed successfully. The probabilities were "high" = 88%, "medium" = 50% and "low" = 12%. Participants were informed that they had twenty minutes to play as many trials as they could, and that three of their win trials would be randomly selected at the end of the task as a monetary reward that they would receive. *Data analysis:* Motivational behavior was operationalized as choosing the hard task (1 - choose, 0 - avoid). Based on previous work, we focused on the tendency for choosing hard task for high rewards (EEfRT - High), which was shown in previous work to correlate with levels of anhedonia[6]. The determination of high and low reward criteria was calculated individually for each participant, based on a dichotomy established by segregating rewards into the upper 30% percentile as 'high' and the lower 70% percentile as 'low' (study 2: 3.5 CAD; study 3: 7 ILS, following [6]). Participants' decisions reflect individual differences in the willingness to expend effort for a reward and represents incentive motivation and anhedonia levels[6]. Poor performance in this task was defined as a frequency of more than 33% of deleted events (response time faster than 200 msec), and led to exclusion from the analysis. To account for differences in number of trials determined by the proportion of hard task choices, we analyzed the first 55 trials of all participants, and discarded the rest. Note that, due to technical issues, the maximal monetary amount in the EEfRT task in study 3 was higher than indicated in the instructions (between 4.3 and 14.8 NIS) for three participants in the first training session (NF group: n=2, sham group: n=1) and three participants in the second session (NF group: n=3). These data were excluded from the analysis.

*Statistical analysis:* To examine how training affected hard task choice behavior for high monetary gains as a function of different choice probabilities, we fitted a GLMM with a binomial logistic regression using the glmer function. The dependent variable was Choice, which was modeled as a binary outcome per trial, participant, and meeting using a logit link function. We modeled the main effects and interaction between group assignment (Test or Sham), Meeting (pre- vs. post-training), and gain Probability in case of success (12%, 50% or 88%), while accounting for the time of presentation (i.e., trial number) as a fixed effect and the individual variability in choice behavior as a random intercept. A Type III ANOVA was performed on the GLMM using Wald chi-square tests to assess fixed effects and interactions.

Probabilistic Selection Task (PST). This task was used as a measure of sensitivity to reward and punishment in reinforcement learning. The task provides indices of learning from positive or negative feedback[7,8]. The PST involves a learning phase and a test phase. In the learning phase three different stimulus pairs are presented to the participants in random order. At each trial, they are presented with two Japanese Hiragana symbols simultaneously on the screen and are required to select one of them. Participants were instructed to select which of the characters is correct by pressing either the left or right button on a keyboard. Each choice is followed by feedback indicating whether the choice was correct or incorrect, but this feedback is probabilistic. The probability of a character to be correct for a given pair varies over the three pairs (AB pair 80/20, CD pair 70/30, EF pair 60/40). Over trials, individuals should learn that the A, C, and E characters are more often associated with positive feedback than the B, D, and F characters. Participants performed the learning blocks (60 trials each) until they met the learning criterion (choosing A over B on 65% of the trials, choosing C over D on 60% of the trials, and choosing E over F on 50% of the trials), to ensure that all participants were at the same performance level before the test phase. In the test phase (90 trials), characters were presented in novel pairs (e.g., AD or BC) at a random order. Participants were instructed to select the "better" character (the one more often associated with positive feedback in the learning phase) and no feedback was provided. To evaluate whether participants learned more from positive or negative feedback we evaluated the accuracy of choosing the A character and avoiding B character. Because stimulus A was the most associated with a positive feedback during training, the more the participant chose stimulus A, when coupled with other symbols, the more the participant learned from reward. Similarly, stimulus B was most often associated with negative feedback during training, such that the more the participant avoided stimulus B, the more the participant learned to avoid the negative feedback (i.e., punishment)[7]. To minimize potential learning effects across meetings - a second version was created using a different set of symbols. The order in which the versions were presented was counterbalanced across subjects. *Data analysis*: The analysis focused on the test phase, where participants' accuracy was examined per meeting and reward type. Accuracy in learning from reward was assessed by indicating, using a binary measure (0 and 1), whether the participant selected the A stimulus in novel pairs (AC, AD, AE, and AF). Similarly, accuracy in learning from punishment was assessed by indicating, also using a binary measure, whether the B stimulus was successfully avoided in novel pairs (BC, BD, BE, and BF). Trial with no response or with a response faster than 200 msec or slower than 4 sec were excluded. Participants who did not learn the AB contingency (chose A over B less than 50% of the trials[7]), and those who had more than 33% of invalid trials were excluded from the analysis. *Statistical analysis*: To examine how the training affected learning from reward and punishments, a GLMM with a binomial logit link was used to separately analyze accuracy in choosing A or avoiding B. The model included group (Test vs. Sham), meeting (post- vs. pre-training) and their interaction, with subject and symbol-pair (i.e., AC, AD, BC, etc.), as random intercepts. To further account for additional factors of no interest that may affect accuracy, the task version and trial order were also included as fixed effects. Model estimation was performed using maximum likelihood with the Laplace approximation, and Type III Wald chi-square tests were used to assess significance.

### Supplementary Results

Supplementary Analysis 2. VS-EFP-NF functional training efficacy (effect on reward function);

Additional Behavioral Tasks - Probabilistic Selection Task and Effort Expenditure for Rewards Task

Study 2: proof of concept: We utilized several computerized tasks that probe reward behavior to assess the possible effects of training. We tested the effects of training on reinforcement learning, by looking at the responses to probabilistic selection task [Frank, 2004] - a task of probabilistic learning, which allows disentangling between patterns of learning from rewards or avoiding punishment. Learning from rewards or punishment was assessed as the accuracy of selecting an often-rewarded symbol (A) or avoiding a seldom-rewarded symbol (B), respectively per trial. Figure S1 depicts this behavior per learning type and group.

Figure S4. Study 2

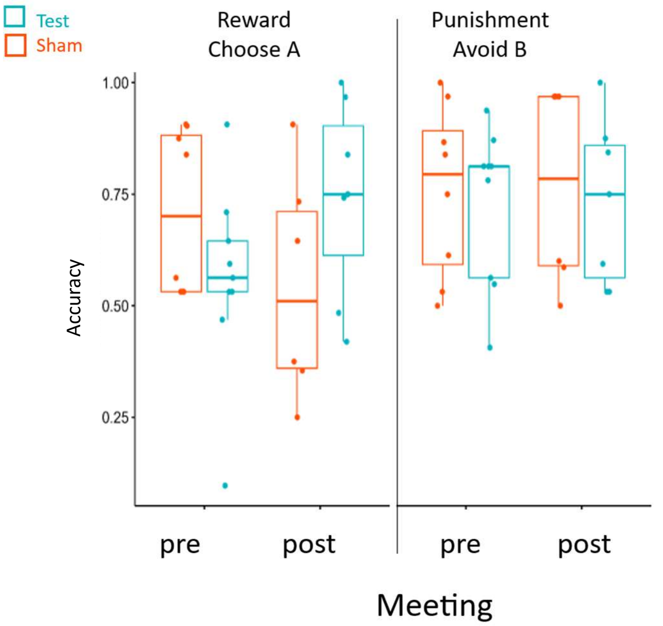


**Figure S1. Study 2, results for the Probabilistic Selection task.** Accuracy rates for learning from reward (left panel) and punishment (right panel) per group (test, sham) and assessment meeting (pre- vs post-training).

Choice accuracy of selecting an often rewarded symbol (A; learning from rewards) was analyzed using a binomial mixed-effects model with fixed effects of Group (test, sham), Meeting (pre, post), and their interaction, as well as nuisance covariates (version and trial order), and a random intercept for participants and symbol pair type. A significant Group × Meeting interaction was observed ($\chi^2(1) = 36.66$, $p < .001$), indicating differential learning across groups. Specifically, the odds of making a correct choice of often rewarded symbols (i.e., learning from rewards) increased more from the pre-training to the post-training meeting in the test group than in the sham group (*Odds-Ratio* = 1.73, 95% CI [1.45, 2.07]; Figure S1). Neither the main effect of Group (*Odds-Ratio* = 0.90, $p = .68$) nor Meeting (*Odds-Ratio* = 0.98, $p = .84$) was significant. A significant effect of task version was observed (*Odds-Ratio* = 1.48, $p < .001$), whereas trial order was not. Together, these results indicated that the neurofeedback training selectively enhanced reward-based learning in the test group. No such interaction effect was found for the accuracy of learning from punishments (avoiding the non-rewarded symbol, B; *Odds-Ratio* = 0.96, 95% CI [0.8, 1.14], $\chi^2(1) = 0.24$, $p = .62$), suggesting that the neurofeedback effect in this study is more relevant to learning from rewards.

Table S1a. Study 2 - Accuracy in learning from rewards (choosing A)

| | Accuracy of Choosing A (yes/no) | | | | |
|---|---|---|---|---|---|
| *Predictors* | *Odds Ratios* | *CI* | *Statistic* | *p* | *df* |
| (Intercept) | 2.40 | 0.97 – 5.95 | 1.89 | 0.059 | Inf |
| Group (test) | 0.90 | 0.53 – 1.51 | -0.41 | 0.682 | Inf |
| Meeting Number | 0.98 | 0.82 – 1.17 | -0.21 | 0.836 | Inf |
| PST version (2) | 1.48 *** | 1.24 – 1.77 | 4.38 | <0.001 | Inf |
| Trial number (z) | 1.04 | 0.88 – 1.23 | 0.45 | 0.651 | Inf |
| Group × Meeting Number | 1.73 *** | 1.45 – 2.07 | 6.05 | <0.001 | Inf |
| **Random Effects** | | | | | |
| $\sigma^2$ | 3.29 | | | | |
| $\tau_{00\ subj}$ | 1.07 | | | | |
| $\tau_{00\ symbol\ pair}$ | 1.14 | | | | |
| ICC | 0.40 | | | | |
| $N_{subj}$ | 17 | | | | |
| $N_{symbol\ pair}$ | 8 | | | | |
| Observations | 945 | | | | |
| Marginal $R^2$ / Conditional $R^2$ | 0.081 / 0.451 | | | | |

** p<0.05 ** p<0.01 *** p<0.001*

Table S1b. Study 2 - Accuracy in avoiding punishments (avoiding B)

| | Accuracy of Avoiding B (yes/no) | | | | |
|---|---|---|---|---|---|
| *Predictors* | *Odds Ratios* | *CI* | *Statistic* | *p* | *df* |
| (Intercept) | 4.75 *** | 2.02 – 11.15 | 3.57 | <0.001 | Inf |
| Group (test) | 1.31 | 0.76 – 2.28 | 0.97 | 0.334 | Inf |
| Meeting Number | 0.88 | 0.74 – 1.06 | -1.34 | 0.179 | Inf |
| PST version (2) | 0.65 *** | 0.54 – 0.78 | -4.67 | <0.001 | Inf |

| | | | | | |
|---|---|---|---|---|---|
| Trial number (z) | 0.96 | 0.81 – 1.12 | -0.55 | 0.584 | Inf |
| Group × Meeting Number | 0.96 | 0.80 – 1.14 | -0.49 | 0.625 | Inf |
| **Random Effects** | | | | | |
| $\sigma^2$ | 3.29 | | | | |
| $\tau_{00\ subj}$ | 1.16 | | | | |
| $\tau_{00\ symbol\ pair}$ | 0.87 | | | | |
| ICC | 0.38 | | | | |
| $N_{subj}$ | 17 | | | | |
| $N_{symbol\ pair}$ | 8 | | | | |
| Observations | 948 | | | | |
| Marginal $R^2$ / Conditional $R^2$ | 0.049 / 0.411 | | | | |

** p<0.05   ** p<0.01   *** p<0.001*

EEfRT task. We next turned to examine another aspect of reward-behavior that relates to incentive motivation - which was assessed using an effort based decision-making task - the EEfRT task. This task assesses willingness to perform a hard task vs. an easy-motor task for gaining rewards of various magnitudes with varying probabilities of gaining. This motivation-related behavior of choosing the hard vs easy task was assessed for the trials offering a chance of earning high monetary gain (threshold set at the 70$^{th}$ percentile, mean threshold = 3.5 ± 0.1 $, following [6]). A logistic generalized mixed-effects model predicting task choice (hard vs. easy) was fitted with group (test vs. sham), meeting (pre vs. post), probability (0.12, 0.5, 0.88), and trial number as predictors, including subject as a random effect. Confirming previous work[6], a significant positive linear effect of probability on hard choice selection was observed (*Odds-Ratio* = 1357.43, 95% CI [414.27, 4447.84], $\chi 2(1)= 141.9$, $p < .001$), indicating an increased likelihood of choosing the hard task with higher gain probabilities. The main effects of group ($p$=0.38) and meeting number ($p$=0.99) were non-significant. The effect of trial number was significant and negative (*Odds-Ratio* =-0.99, 95% CI [0.98,1.00], $\chi 2(1)= 3.88$, $p=0.05$), suggesting lower probability of choosing the hard task in later trials. Importantly, significant interaction effects were observed. The group × meeting interaction (*Odds-Ratio* = 2.4, 95% CI [1.32, 4.38], $\chi 2(1)= 8.24$, $p=0.004$; Figure S2) was significant, indicating that the group effect varied by meeting, being more pronounced following training, with greater overall tendency to opt for hard choices among test relative to sham following training (post hoc test: post-training: Odds-Ratio (sham / test) = 0.13, *z*.ratio = -2, $p = 0.04$; pre-training: Odds-Ratio (sham / test) = 0.52, *z*.ratio = -0.65, $p = 0.52$). The group x meeting x probability interaction was also significant (Odds-ratio = 0.34, 95% CI [0.13, 0.89], $\chi 2(1)= 4.88$, $p=0.03$), indicating that the effect of NF training on incentive behavior following training negatively varied as a function of reward probabilities, being more pronounced for lower probabilities (12% post-training: odds Ratio (sham / test) = 0.075, *z*.ratio = -2.15, $p = 0.03$; 50% post-training: odds Ratio (sham / test) = 0.13, *z*.ratio = -2.01, $p = 0.04$; 88% post-training: odds Ratio (sham / test) = 0.24, *z*.ratio = -1.32, $p = 0.18$). Together, this analysis suggests that following VS-EFP NF but not sham NF, a greater incentive motivation behavior is evident, especially under lower reward probabilities.

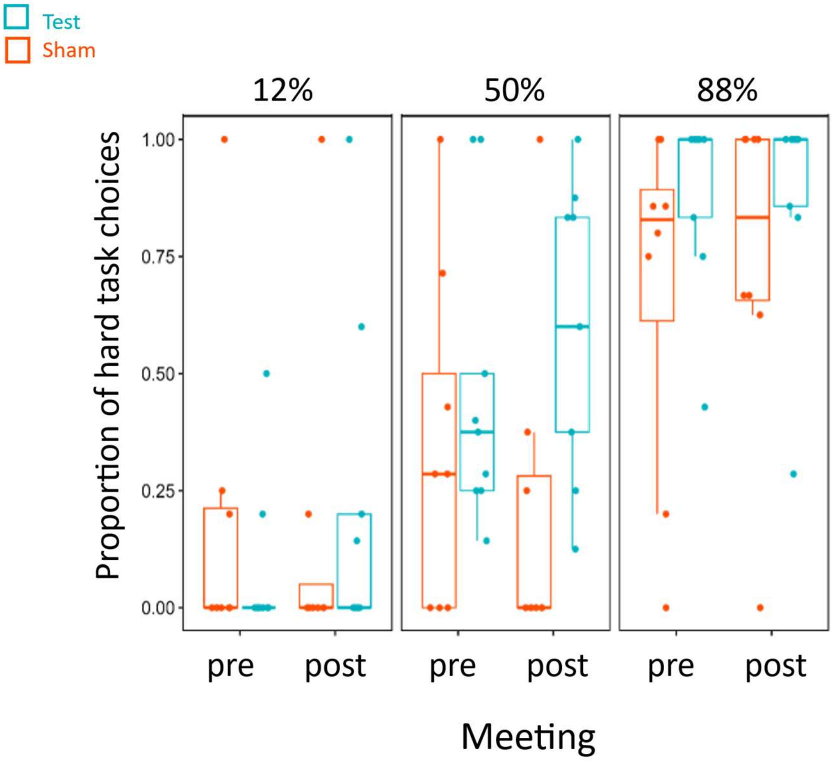


**Figure S2. Study 2, Incentive motivation behavior. Results for the EEfRT task**. Rates of choosing the hard task for different gain probabilities (12%, 50% and 88%) per group (test, sham) and meeting (pre- vs post-training)

Table S2. EEfRT task: tendency of selecting hard choices for a high monetary reward – study 2

| | Choice | | | | |
|---|---|---|---|---|---|
| *Predictors* | *Odds Ratios* | *CI* | *Statistic* | *p* | *df* |
| (Intercept) | 0.02 *** | 0.01 – 0.08 | -5.96 | <0.001 | Inf |
| Group (test) | 0.60 | 0.19 – 1.88 | -0.88 | 0.379 | Inf |
| Meeting (post) | 1.00 | 0.55 – 1.81 | -0.01 | 0.993 | Inf |
| Probability | 1357.43 *** | 414.27 – 4447.84 | 11.91 | <0.001 | Inf |
| Trial number (time) | 0.99 * | 0.98 – 1.00 | -1.97 | 0.049 | Inf |
| Group (test) × Meeting (post) | 2.40 ** | 1.32 – 4.38 | 2.87 | 0.004 | Inf |

| | | | | | |
|---|---|---|---|---|---|
| Group (test) × probability | 0.74 | 0.24 – 2.29 | -0.53 | 0.596 | Inf |
| Meeting (post) × Probability | 1.00 | 0.39 – 2.57 | 0.00 | 0.999 | Inf |
| Group (test) × Probability x Meeting (post) | 0.34 * | 0.13 – 0.89 | -2.21 | **0.027** | Inf |
| **Random Effects** | | | | | |
| $\sigma^2$ | 3.29 | | | | |
| $\tau_{00\ subj}$ | 3.83 | | | | |
| ICC | 0.54 | | | | |
| $N_{subj}$ | 18 | | | | |
| Observations | 657 | | | | |
| Marginal $R^2$ / Conditional $R^2$ | 0.449 / 0.745 | | | | |

** p<0.05   ** p<0.01   *** p<0.001*

Study 3: Validation in an independent cohort (N=40):

PST. To examine whether the finding from study 2, indicating an influence of training on learning from rewards, we ran a similar logistic generalized mixed-effects model predicting accuracy in choosing A. A main effect of Meeting was found (Odds-Ratio= 1.21, 95% CI [1. 08, 1.36], $\chi^2(1)$ = 10.24, $p$ = .001;  Figure S3), indicating a decrease in accuracy across baseline and outcome meetings (*pre vs. post*: Odds-Ratio = 1.47, $p$ = .001), along with an effect of version (Odds Ratio = 2.11, 95% CI [1.88, 2.38] , $\chi^2(1)$ = 152.4, $p$ < .001). This pattern is consistent with the findings of Study 2. However, no interaction between group and meeting was found (Odds-Ratio = 1.02, 95% CI [0.91, 1.14], $\chi^2(1)$ = 0.07, $p$ = .79), failing to replicate the previously observed effect of VS-EFP-NF training on the pattern of learning from rewards.

Table S3. Study 3 - Accuracy in learning from rewards (choose A)

| | Accuracy of Choosing A (yes/no) |
|---|---|

| *Predictors* | *Odds Ratios* | *CI* | *Statistic* | *p* | *df* |
|---|---|---|---|---|---|
| (Intercept) | 2.95 *** | 1.89 – 4.62 | 4.75 | **<0.001** | Inf |
| Group (test) | 0.97 | 0.72 – 1.30 | -0.21 | 0.832 | Inf |
| Meeting Number | 1.21 ** | 1.08 – 1.36 | 3.20 | **0.001** | Inf |
| PST version (2) | 2.11 *** | 1.88 – 2.38 | 12.35 | **<0.001** | Inf |
| Trial number (z) | 1.00 | 0.90 – 1.11 | -0.07 | 0.940 | Inf |
| Group × Meeting Number | 1.02 | 0.91 – 1.14 | 0.26 | 0.793 | Inf |
| **Random Effects** | | | | | |
| $\sigma^2$ | 3.29 | | | | |
| $\tau_{00\ subj}$ | 0.74 | | | | |
| $\tau_{00\ symbol\ pair}$ | 0.24 | | | | |
| ICC | 0.23 | | | | |
| $N_{subj}$ | 40 | | | | |
| $N_{symbol\ pair}$ | 8 | | | | |
| Observations | 2200 | | | | |
| Marginal $R^2$ / Conditional $R^2$ | 0.115 / 0.318 | | | | |

** p<0.05   ** p<0.01   *** p<0.001*

Figure S6. Study 3

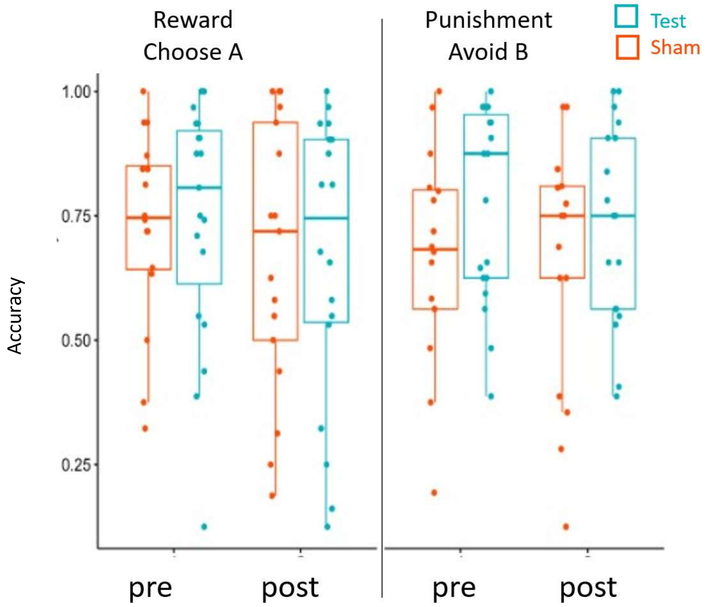


Figure S3. Study 3, results for the PST task: accuracy rates for learning from reward (left panel) and punishment (right panel) per group (test, sham) and session (pre vs post)

EEfRT task: A logistic generalized mixed-effects model predicting task choice proportion (hard vs. easy) was fitted with group, meeting, probability, and trial number as predictors, including subject as a random effect for all valid datasets (test group: N = 14, sham group: N=19). Replicating study 2 and confirming previous work[6], a significant positive effect of probability on hard choice selection was observed (*Odds-Ratio* = 481.34, 95% CI [230.14, 1006.72], $\chi 2(1)$= 269.16, $p$ < .001; Figure S4), indicating an increased likelihood of choosing the hard task with higher gain probabilities. Yet, in contrary to the findings from study 2, no interaction was found between group and meeting (*Odd-Ratio* = 0.98, 95% CI [0.54, 1.78], $\chi 2(1)$ = 0.01, $p$ = 0.94), and no interaction between group, meeting and probability was evident (*Odds-Ratio* = 0.71, 95% CI [0.39, 1.29], $\chi 2(1)$ = 1.25, $p$ = 0.26).

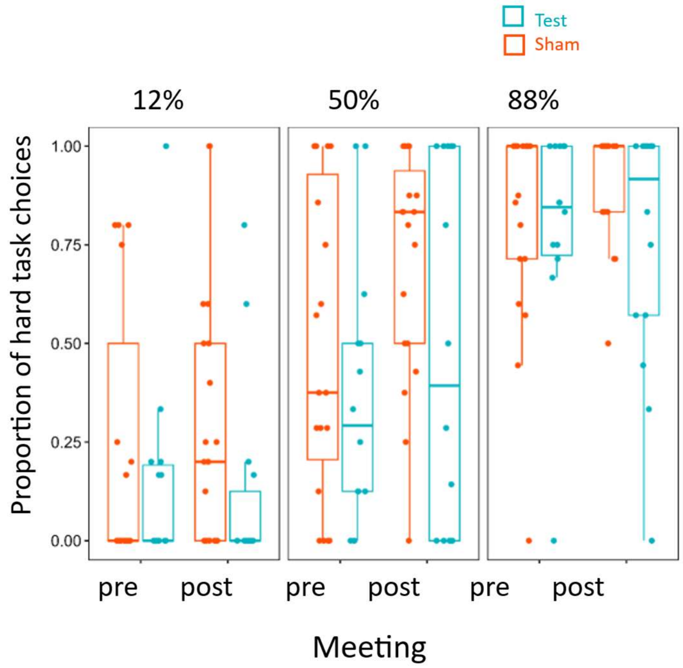


Figure S4. Study 3. Motivation behavior. Results for the EEfRT task: rates of choosing the hard task for different gain probabilities (12%, 50% and 88%) per group (VS-EFP, sham) and session (pre vs post)

Table S4. study 3 - EEfRT task: tendency of selecting hard choices for a high monetary reward

| | Hard Task Choice | | | | |
|---|---|---|---|---|---|
| *Predictors* | *Odds Ratios* | *CI* | *Statistic* | *p* | *df* |
| (Intercept) | 0.06 *** | 0.02 – 0.13 | -6.82 | **<0.001** | Inf |
| Group (test) | 2.26 * | 1.05 – 4.84 | 2.10 | **0.036** | Inf |
| Meeting (post) | 0.75 | 0.52 – 1.07 | -1.59 | 0.112 | Inf |
| Probability | 481.34 *** | 230.14 – 1006.72 | 16.41 | **<0.001** | Inf |
| Trial number (time) | 1.00 | 0.99 – 1.00 | -1.00 | 0.317 | Inf |
| Group (test) × Meeting (post) | 0.98 | 0.69 – 1.40 | -0.11 | 0.914 | Inf |
| Group (test) × probability | 0.66 | 0.32 – 1.35 | -1.13 | 0.258 | Inf |
| Meeting (post) × Probability | 0.98 | 0.54 – 1.78 | -0.08 | 0.936 | Inf |

| | | | | | |
|---|---|---|---|---|---|
| Group (test) × Probability x Meeting (post) | 0.71 | 0.39 – 1.29 | -1.12 | 0.263 | Inf |
| **Random Effects** | | | | | |
| $\sigma^2$ | 3.29 | | | | |
| $\tau_{00\ subj}$ | 3.44 | | | | |
| ICC | 0.51 | | | | |
| $N_{subj}$ | 33 | | | | |
| Observations | 1280 | | | | |
| Marginal $R^2$ / Conditional $R^2$ | 0.379 / 0.697 | | | | |

** p<0.05 ** p<0.01 *** p<0.001*
